# Molecular hallmarks of excitatory and inhibitory neuronal resilience and resistance to Alzheimer’s disease

**DOI:** 10.1101/2025.01.13.632801

**Authors:** Isabel Castanho, Pourya Naderi Yeganeh, Carles A. Boix, Sarah L. Morgan, Hansruedi Mathys, Dmitry Prokopenko, Bartholomew White, Larisa M. Soto, Giulia Pegoraro, Saloni Shah, Athanasios Ploumakis, Nikolas Kalavros, David A. Bennett, Christoph Lange, Doo Yeon Kim, Lars Bertram, Li-Huei Tsai, Manolis Kellis, Rudolph E. Tanzi, Winston Hide

**Affiliations:** Harvard Medical School, Boston, MA, USA; Department of Pathology, Beth Israel Deaconess Medical Center, Boston, MA, USA; Computer Science and Artificial Intelligence Laboratory, MIT, Cambridge, MA 02139, USA; Broad Institute of MIT and Harvard, Cambridge, MA 02142, USA; Centre for Neuroscience, Surgery and Trauma, Blizard Institute, Queen Mary University of London, London E1 2AT, UK; University of Pittsburgh Brain Institute, University of Pittsburgh School of Medicine, Pittsburgh, PA 15261, USA; Picower Institute for Learning and Memory, MIT, Cambridge, MA 02139, USA; Genetics and Aging Research Unit, The Henry and Allison McCance Center for Brain Health, Department of Neurology, Massachusetts General Hospital, Boston, MA, United States; Medical School, University of Exeter, Exeter EX2 5DW, UK; Spatial Technologies Unit, Beth Israel Deaconess Medical Center, Boston, MA, USA; Rush Alzheimer’s Disease Center, Rush University Medical Center, 1750 W Harrison Street, Suite 1000, Chicago, IL, 60612, USA; Department of Biostatistics, Harvard T.H. Chan School of Public Health, 677 Huntington Ave, 02115, Boston, MA, USA; Lübeck Interdisciplinary Platform for Genome Analytics, Institutes of Neurogenetics and Cardiogenetics, University of Lübeck, Lübeck, Germany; Department of Psychology, University of Oslo, Oslo, Norway; Department of Brain and Cognitive Sciences, MIT, Cambridge, MA 02139, USA

**Author notes:** These authors contributed equally to this work (Isabel Castanho, Pourya Naderi Yeganeh). Correspondence to: Winston Hide.

**Keywords:** Cognitive resilience, Alzheimer’s disease, gene expression, transcriptomics, vulnerability, genetics, rare variants

## Abstract

**Background:** A significant proportion of individuals maintain healthy cognitive function despite having extensive Alzheimer’s disease (AD) pathology, known as cognitive resilience. Understanding the molecular mechanisms that protect these individuals can identify therapeutic targets for AD dementia. This study aims to define molecular and cellular signatures of cognitive resilience, protection and resistance, by integrating genetics, bulk RNA, and single-nucleus RNA sequencing data across multiple brain regions from AD, resilient, and control individuals.

**Methods:** We analyzed data from the Religious Order Study and the Rush Memory and Aging Project (ROSMAP), including bulk (n=631) and multi-regional single nucleus (n=48) RNA sequencing. Subjects were categorized into AD, resilient, and control based on β-amyloid and tau pathology, and cognitive status. We identified and prioritized protected cell populations using whole genome sequencing-derived genetic variants, transcriptomic profiling, and cellular composition distribution.

**Results:** Transcriptomic results, supported by GWAS-derived polygenic risk scores, place cognitive resilience as an intermediate state in the AD continuum. Tissue-level analysis revealed 43 genes enriched in nucleic acid metabolism and signaling that were differentially expressed between AD and resilience. Only GFAP (upregulated) and KLF4 (downregulated) showed differential expression in resilience compared to controls. Cellular resilience involved reorganization of protein folding and degradation pathways, with downregulation of Hsp90 and selective upregulation of Hsp40, Hsp70, and Hsp110 families in excitatory neurons. Excitatory neuronal subpopulations in the entorhinal cortex (ATP8B1+ and MEF2C^high^) exhibited unique resilience signaling through neurotrophin (modulated by LINGO1) and angiopoietin (ANGPT2/TEK) pathways. We identified MEF2C, ATP8B1, and RELN as key markers of resilient excitatory neuronal populations, characterized by selective vulnerability in AD. Protective rare variant enrichment highlighted vulnerable populations, including somatostatin (SST) inhibitory interneurons, validated through immunofluorescence showing co-expression of rare variant associated RBFOX1 and KIF26B in SST+ neurons in the dorsolateral prefrontal cortex. The maintenance of excitatory-inhibitory balance emerges as a key characteristic of resilience.

**Conclusions:** We identified molecular and cellular hallmarks of cognitive resilience, an intermediate state in the AD continuum. Resilience mechanisms include preservation of neuronal function, maintenance of excitatory/inhibitory balance, and activation of protective signaling pathways. Specific excitatory neuronal populations appear to play a central role in mediating cognitive resilience, while a subset of vulnerable SST interneurons likely provide compensation against AD-associated dysregulation. This study offers a framework to leverage natural protective mechanisms to mitigate neurodegeneration and preserve cognition in AD.

## Background

The search for effective disease-modifying treatments for Alzheimer’s disease (AD) has primarily focused on targeting β-amyloid (Aβ) with limited success. Historically, a definitive AD diagnosis has required postmortem observation of Aβ and tau neuropathology. However, some individuals maintain healthy cognitive function despite meeting neuropathological criteria for AD at autopsy, termed as *cognitive resilience* (or cognitive reserve) [1,2]. In contrast, AD resistance describes individuals who do not develop AD neuropathology or cognitive decline, even in the presence of the strongest AD risk factor: advanced aging (**Figure 1A**). Understanding natural protective mechanisms is a potent approach to developing novel AD therapeutics beyond Aβ targeting. However, the precise molecular systems of protection against AD dementia via resilience or resistance and their temporal relationship to AD pathogenesis remain poorly understood.

**Figure 1.**
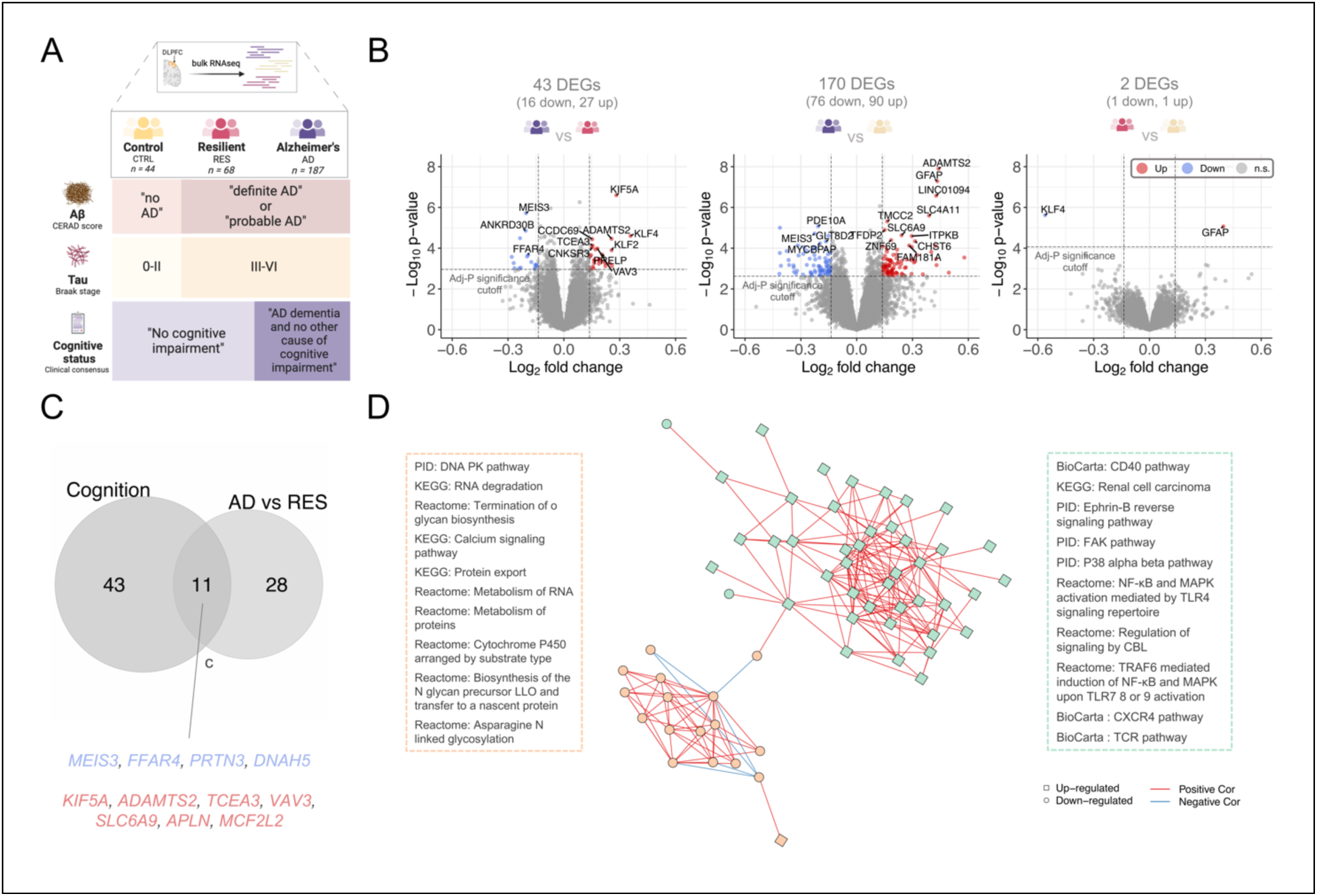
Transcriptomic and pathway signatures of cognitive resilience against AD pathology. **(A)** Overview of study design. Using levels of Aβ plaques and neurofibrillary tangles, and presence/absence of cognitive impairment, ROSMAP donors were classified into three major categories: Control (CTRL, CERAD “no AD”, Braak 0-II, and consensus cognitive diagnosis “no cognitive impairment”), Resilient (RES, CERAD “definite AD” or “probable AD”, Braak III-VI, and consensus cognitive diagnosis “no cognitive impairment”), and AD (CERAD “definite AD” or “probable AD”, Braak III-VI, and consensus cognitive diagnosis “Alzheimer’s dementia and no other cause of cognitive impairment”). **(B)** Volcano plots showing significantly (adj-P < 0.1) differentially expressed genes (DEGs) in AD compared to resilient individuals (ADvsRES), AD compared to controls (ADvsCTRL), and resilient compared to controls (RESvsCTRL). DEGs with log2FC < -log2(1.1) are highlighted in blue, and DEGs with log2FC > log2(1.1) are highlighted in red. The horizontal lines represent FDR-adjusted P-value = 0.1. **(C)** Venn diagram showing common genes identified as DEGs in ADvsRES and identified as associated with cognitive decline. Down-regulated genes are shown in blue, and up-regulated genes in red. **(D)** Two major classes of dysregulated functions in ADvsRES identified by pathway activity analysis. Nodes represent pathways with significant dysregulation of activity (q-value < 0.1) in ADvsRES. Out of a total of 99 dysregulated pathways between ADvsRES, 62 organized in two unsupervised clusters of expression. Node shapes denote up-regulation and down-regulation in AD. Edges represent co-expression of pathways based on the Pathway Co-expression Network background. Pathway activity profiles were determined using the PanomiR software package. Pathway dysregulation p-values were determined using the Limma package’s linear regression models contrasting between ADvsRES and accounting for confounding covariates such as age, batch, and RNA integrity number. Figure created in part with BioRender.com.

Genetic and high-throughput studies have identified molecular features associated with AD protection and risk [3–8]. These features are often attributed to resistance or vulnerability. Genome-wide association studies (GWAS) have highlighted prominent genetic factors such as *APOE* e4 and *TREM2*, which explain differences between AD and healthy individuals[4] and are described as protection/risk mechanisms. Most healthy populations in these studies lack AD pathology and are not stratified by resilience phenotypes[9,10]. It remains unclear whether protection occurs at the level of AD pathology or AD dementia.

Understanding how certain individuals prevent cognitive decline despite exposure to advanced aging can lead to effective approaches tailored for specific stages of the disease. Recent genetic studies have shown that specific mutations in *APOE* and *ATP8B1*, along with sex-linked loci, are associated with resilience [3–5,11]. MEF2C (myocyte enhancer factor 2C) promotes resilience in both humans and mouse models [12]. Strong evidence supports MEF2C’s role in hyperexcitability of excitatory neurons [12]. These findings underscore the importance of understanding gene signatures in their cellular contexts to elucidate disease mechanisms. Single-cell genomics has revealed disease-associated vulnerability in specific cell types such as somatostatin GABAergic inhibitory neurons, and distinct gene regulatory programs differentiating vulnerable individuals [6,13–18]. Recent multi-region single-nuclei transcriptomic analysis has found specific RELN-expressing excitatory neuronal populations in the entorhinal cortex with selective vulnerability to AD progression. Using a continuous measure of resilience, the same study has found predominance of enrichment of resilience-associated genes in astrocytes [18]. Maintaining cellular populations, such as SST inhibitory neurons or MEF2C and RELN excitatory neurons, appears to be integral to cognitive protection against AD. Contextualizing these protective mechanisms in terms of resilience, resistance, and the temporal order of AD pathogenesis is critical for deriving translational insights. Since natural protective functions against AD pathogenesis are dynamic, strategies to leverage them will depend on the individual’s pathological status.

Resilience acts as a safeguard for cognition when resistance mechanisms fail to prevent pathological lesions. Disentangling the factors driving resilience from those underlying resistance is crucial to combat AD in individuals with or at risk of neuropathology. The difference between molecular and cellular underpinnings of resilience and resistance remain insufficiently described. In this study, we systematically investigate molecular and cellular determinants of AD resilience. Leveraging previous work that used a continuous measure of resilience [18], we have now applied a more discrete model to categorize resilience in the aging brain, paving the way for targeted interventions that could preserve cognitive function.

## Results

We employed integrative transcriptomic and single-cell analyses to explore the molecular and cellular mechanisms underlying resilience. We performed molecular characterization across transcriptomes, assessed neuronal subpopulations for their roles in resilience, and predicted resilience intercellular communication dynamics. We employed a genetic enrichment approach to determine if populations of cells were likely to be vulnerable or associated with protection.

### Resilience as an intermediate transcriptomic state in the AD continuum

To define the molecular and functional signatures of resilience and to stratify them from those of AD and healthy individuals, we used bulk RNAseq data from the dorsolateral prefrontal cortex (DLPFC) of individuals from the Religious Order Study and Memory and Aging Project cohort (ROSMAP, n = 631) [19]. We classified the ROSMAP subject into mutually exclusive groups of resilience, AD, and healthy controls using levels of Aβ plaques and neurofibrillary tangles and presence/absence of cognitive impairment [20] (**Figure 1A**, **Methods,** phenotypic characteristics per group are shown in **Figure S2**).

We found 43 differentially expressed genes (DEGs) in AD compared to resilient (adjusted P < 0.1 and |log2FC| > log(1.1)) (ADvsRES, **Figure 1B, Table S1**), and 170 DEGs in ADvsCTRL (**Table S2**). In marked contrast, only two genes, *GFAP* (encoding for glial fibrillary acidic protein) and *KLF4* (KLF transcription factor 4), reached significance for differential expression in resilient subjects compared to control (RESvsCTRL, **Table S3**). *GFAP* was one of our top DEGs in ADvsCTRL (**Table S2**, log2FC = 0.43, adjusted P = 4.0 x 10^-4^). On the other hand, *KLF4* exhibited an opposing expression pattern between ADvsRES (Log2FC = 0.36, adjusted P = 0.035) compared to RESvsCTRL (Log2FC = −0.56, adjusted P = 0.038). This suggests that KLF4 has resilience-specific down-regulation.

To further charachterize resilience, we included a presympatomic category (PRE) in our analysis, made up of individuals with with mild cognitive impairment and advanced AD pathology (see **Methods**). A comparison of presymptomatic versus resilient subjects yield no DEGs, whereas there were 68 DEGs between presymptomatic and AD (**Table S4**). These results indicate that the majority of the transcriptomic changes in the AD continuum emerge during the transition from a resilient/presymptomatic state to late-stage AD dementia, rather than at the onset of pathology without cognitive impairment.

The major difference between the AD group with the presymptomatic and resilience groups was the presence/absence of dementia. To determine the extent to which resilience signatures are explained by signatures of cognitive loss, we compared resilience-associated and cognitive-associated genes to identify common signatures. Cognitive-associated genes were identified using ordinal categorical regression (proportional odds model) irrespective of group classification and adjusted for the burden of pathology. We identified 54 genes significantly (adj-P < 0.1) associated with cognitive decline when adjusted for pathology burden (**Table S5**). Of the 54 cognitive impairment-associated genes, 11 (20.4%) were also DEG between AD and resilience (**Figure 1C** and **Table S5**). Our results show that a considerable portion of resilience-associated genes are directly correlated with the events that lead to cognitive and neuronal loss in dementia.

To establish whether resilience is an intermediate phenotype or a distinct trajectory from AD pathogenesis, we compared the genetic risk profile of resilient subjects with AD. We used AD polygenic risk scores (AD-PRS), which evaluate the risk prediction for AD considering AD-associated single nucleotide polymorphisms (SNPs). Across all subjects with whole genome sequencing (WGS) profiles from the ROSMAP cohort [20] resilient subjects exhibited an intermediate genetic risk profile score, significantly different from both AD and controls (**Figure S1B**). Combining transcriptomic and genetic evidence, our analysis suggests that resilience is the ability to maintain a presymptomatic state that falls within the spectrum of AD pathogenesis.

### Functional characterization of resilience transcriptomes

To functionally characterize pathway dysregulation events, we used the differences in transcriptomics in AD compared to resilient subjects. We used pathway activity analysis, part of the *PanomiR* package [21,22], to perform activity summarization to represent the *overall* activity of genes within known pathways. Applied to ROSMAP RNAseq data and using 1329 background pathways from the MSigDB database, we identified significant dysregulation of 99 pathways (q-value < 0.1) in ADvsRES (**Table S6**). Leveraging an independent canonical co-expression map of the pathways and unsupervised clustering, we identified two super-groups of pathways that dominated the differences between AD and resilience subjects. The largest cluster included 43 dysregulated pathways related to up-regulation of signaling in AD, including NFKB (nuclear factor kappa-light-chain-enhancer) activation, MAPK (mitogen-activated protein kinases) activation, and FAK (focal adhesion kinase) signaling. The second largest cluster included 19 dysregulated pathways representing down-regulation of RNA machinery and metabolism in AD compared to resilience, including RNA degradation and RNA metabolism, calcium signaling, and protein metabolism and export. Comparison of AD with control and presymptomatic groups found 51 dysregulated pathways (q-value < 0.1) in ADvsCTRL (**Table S8**) and 290 dysregulated pathways (q-value < 0.1) in ADvsPRE (**Table S10**). Concordant with gene-level analysis, no significant changes were identified between resilience and presymptomatic groups (**Table S9**). Taken together, pathway activity analysis defines resilience and presymptomatic states with maintenance of RNA/DNA processing pathways lost during transition to AD dementia, and replacement by activation of signaling cascades.

### Ubiquitous markers of cellular resilience

To understand if there were ubiquitous molecular signatures of resilience, we investigated cell-specific signatures of existing single-nucleus RNAseq (snRNAseq) profiles from DLPFC, entorhinal cortex (EC), and hippocampus (HC) from a multi-region dataset generated from the ROSMAP cohort [18] (n = 48). Similar to the bulk tissue study here, we used levels of Aβ plaques and neurofibrillary tangles, and presence/absence of cognitive impairment, to classify ROSMAP subjects into AD, resilient, and control (**Table S11**). Phenotypic distributions per group are shown in **Figures S5-S7**. Major cell types were identified as previously described [18], and cell subtypes were identified from brain region-, cell type-specific subclusters, and annotated using two independent brain references (see **Methods** and **Table S14**).

We performed DEG analysis on individual cell types in the DLPFC. They showed a consistently higher number of dysregulation events in ADvsRES compared to ADvsCTRL (**Figure S8A, Table S12**). This pattern was reflected in the EC and HC. Astrocytes, particularly in the DLPFC, exhibited more pronounced upregulation and downregulation of genes in ADvsRES, with a notable increase in genes downregulated in AD (**Figure S8B**). Additionally, upregulated DEGs showed greater changes in RESvsCTRL when compared to ADvsCTRL. Excitatory neurons from all brain regions also exhibited a higher number of DEGs in ADvsRES compared to ADvsCTRL (**Figure S8B**), which was also the case for inhibitory neurons from the DLPFC and EC (**Figure S8B**).

Concordant with our observations from bulk RNAseq data, the snRNAseq expression of *GFAP* in the DLPFC was significantly down-regulated in AD compared to resilience (ADvsRES) and up-regulated in resilience compared to controls (RESvsCTRL). *GFAP* was identified as the top DEG in DLPFC astrocytes in RESvsCTRL (**Figure 2A**, **Table S13**). No significant differences for *GFAP* were found in ADvsRES in the EC and HC (**Table S13**).

**Figure 2.**
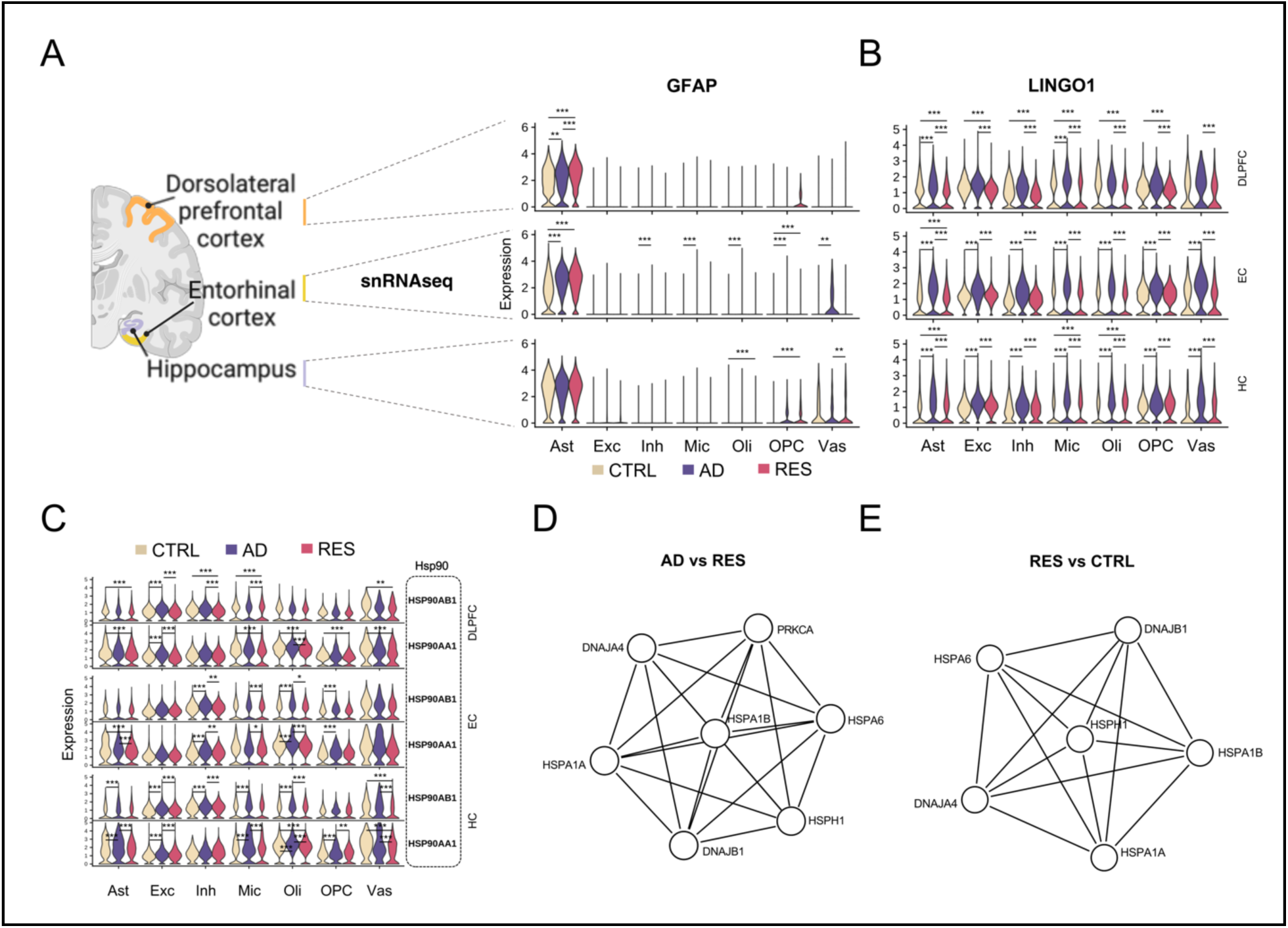
Cell-specific transcriptomic signatures of cognitive resilience against AD pathology. **(A-C)** Violin plots showing gene expression for selected DEGs across different major cell types from each brain region. Log2FC, adjusted P-values (adj-P), and direction of change (first diagnostic group compared to the second group) are shown in **Table S14**. * Adj-P < 0.05, ** Adj-P < 0.01, *** Adj-P < 0.001. **(A)** Changes in the expression of *GFAP*. **(B)** Down-regulation of *LINGO1* in cognitive resilience. **(C)** Down-regulation of Hsp90 (heat shock protein 90) family members in cognitive resilience. **(D-E)** Protein-protein interaction (PPI) networks in excitatory neurons generated using Metascape. Molecular Complex Detection (MCODE) algorithm network clusters (modules) showing the subset of proteins that form physical interactions with at least one other member in the list. The protein networks were constructed based on physical interactions among all input gene lists. **(D)** PPI network clusters detected from genes up-regulated in excitatory neurons (DLPFC) in resilience from ADvsRES. The three best-scoring terms by p-value from pathway and process enrichment analysis for this module were “chaperone cofactor-dependent protein refolding” (GO:0051085, Log10(P) = −12.6), “’de novo’ post-translational protein folding” (GO:0051084, Log10(P) = −12.3), and “’de novo’ protein folding” (GO:0006458, Log10(P) = −12.1). **(E)** PPI network cluster detected from genes up-regulated in excitatory neurons (DLPFC) in resilience from RESvsCTRL. The three best-scoring terms by p-value from pathway and process enrichment analysis for this module were “chaperone cofactor-dependent protein refolding” (GO:0051085, Log10(P) = −13.2), “’de novo’ post-translational protein folding” (GO:0051084, Log10(P) = −12.9), and “’de novo’ protein folding” (GO:0006458, Log10(P) = - 12.6).

LINGO1, the most significant DEG across major cell types in the DLPFC, was consistently upregulated in AD but downregulated in resilient subjects across the three brain regions (**Figure 2B, Table S13**). In the DLPFC, LINGO1 was significantly reduced in resilience (RESvsCTRL) across most cell types, except vascular/epithelial cells, while in the EC, changes were subtle and below the FC cutoff (**Table S13**). In contrast, LINGO1 was upregulated in resilience (RESvsCTRL) in astrocytes, microglia, and oligodendrocytes in the HC, suggesting regional specificity to cortical brain areas.

The expression levels of genes (*HSP90AB1* and *HSP90AA1*) coding for two members of the heat shock protein 90 (Hsp90) family were down-regulated in multiple cell populations in resilience in all brain regions investigated. *HSP90AB1* was significantly up-regulated in AD (ADvsRES) in excitatory and inhibitory neurons, and in oligodendrocytes in the DLPFC; in inhibitory neurons, microglia, and oligodendrocytes in the EC; and excitatory neurons, inhibitory neurons, oligodendrocytes, and vascular/epithelial cells in the HC (**Figure 2C**, **Table S13**).

*HSP90AB1* was also down-regulated in resilience (RESvsCTRL) in the DLPFC in astrocytes, inhibitory neurons, oligodendrocytes, and vascular/epithelial cells. Another Hsp90 member, *HSP90AA1*, was up-regulated in AD (ADvsRES) in excitatory neurons and oligodendrocytes, and down-regulated in resilience (RESvsCTRL) in all major cell types except neurons in the DLPFC (**Figure 2C**, **Table S13**). Furthermore, *HSP90AA1* was significantly down-regulated in resilience (RESvsCTRL) in astrocytes in the EC and oligodendrocytes and vascular and epithelial cells in the HC.

Gene ontology (GO) enrichment analysis for each of the lists of DEGs in major cell types from each comparison (ADvsRES, ADvsCTRL, and RESvsCTRL, with up and down-regulated genes investigated separately) identified multiple terms related to protein folding up-regulated in resilience (**Figure S9**). Protein-protein interaction (PPI) network analysis generated a single cluster from the list of DEGs up-regulated in excitatory neurons in DLPFC tissue from resilient individuals in ADvsRES (**Figure 2D**), which was enriched in terms related to protein folding and included members of the families Hsp40 (*DNAJA4*, *DNAJB1*), Hsp70 (*HSPA1A*, *HSPA1B*, *HSPA6*), and Hsp110 (*HSPH1*). Concurrently, PPI analysis for DEGs up-regulated in excitatory neurons in RESvsCTRL resulted in one cluster enriched for “protein folding” ontology terms (**Figure 2E**), which also included members of the families Hsp40 (*DNAJA4*, *DNAJB1*), Hsp70 (*HSPA1A*, *HSPA1B*, *HSPA6*), and Hsp110 (*HSPH1*). Examination of the differential expression of genes from the Hsp40, Hsp70, and Hsp110 families found a consistent upregulation in neurons in resilience - and downregulation in AD in excitatory neurons - and mostly downregulation in glial cells (**Table S13**).**’**

### Region-specific protection and resistance phenotypes in inhibitory neurons

To define vulnerable and resilient cell types, we charted cellular expression bias of genes associated with protection and vulnerability and contrasted the results with differential abundance across AD, resilient, and aged-matched healthy conditions. To chart the cellular expression bias of genes associated with protection, we investigated the overall expression of rare and common AD variants in individual cell types annotated from the multi-region snRNAseq profiles using Expression Weighted Cell Enrichment (EWCE). Common genetic variants were selected based on a recent study of approximately 800K individuals [9]. Genes associated with rare variants (single nucleotide polymorphisms with minor allele frequency ≤1%), were selected as part of our recent whole-genome survey of families afflicted with AD [23,24]. Rare genetic variants found in asymptomatic family members were classified as ‘protective’, and genetic variants associated with AD were classified as ‘risk’ variants (see Methods and limitations of study).

Inhibitory neurons from the DLPFC and EC (**Figure 3A-B**) were significantly enriched for genes from ‘protective’ variants (**Figure 3C**, **Figure S10C**). This was also true for those in HC (Figure S10C). Genes from ‘risk’ variants were significantly enriched in inhibitory neurons and oligodendrocyte progenitor cells (OPCs) in all brain regions tested (**Figure S10A-C**). We confirmed and validated this trend in a population of inhibitory neurons from the DLPFC in an independent snRNAseq dataset [25](**Figure S10D**). Genes associated with common variants [9], showed a strong bias towards genes expressed in microglia and immune cells in all brain regions (**Figure S10E**), consistent with current reports [26]. Our results suggest that genes that harbor rare genetic variants, which tend to have stronger effect sizes, reveal a previously undescribed bias in inhibitory neurons for genes associated with protection pathways[27].

**Figure 3.**
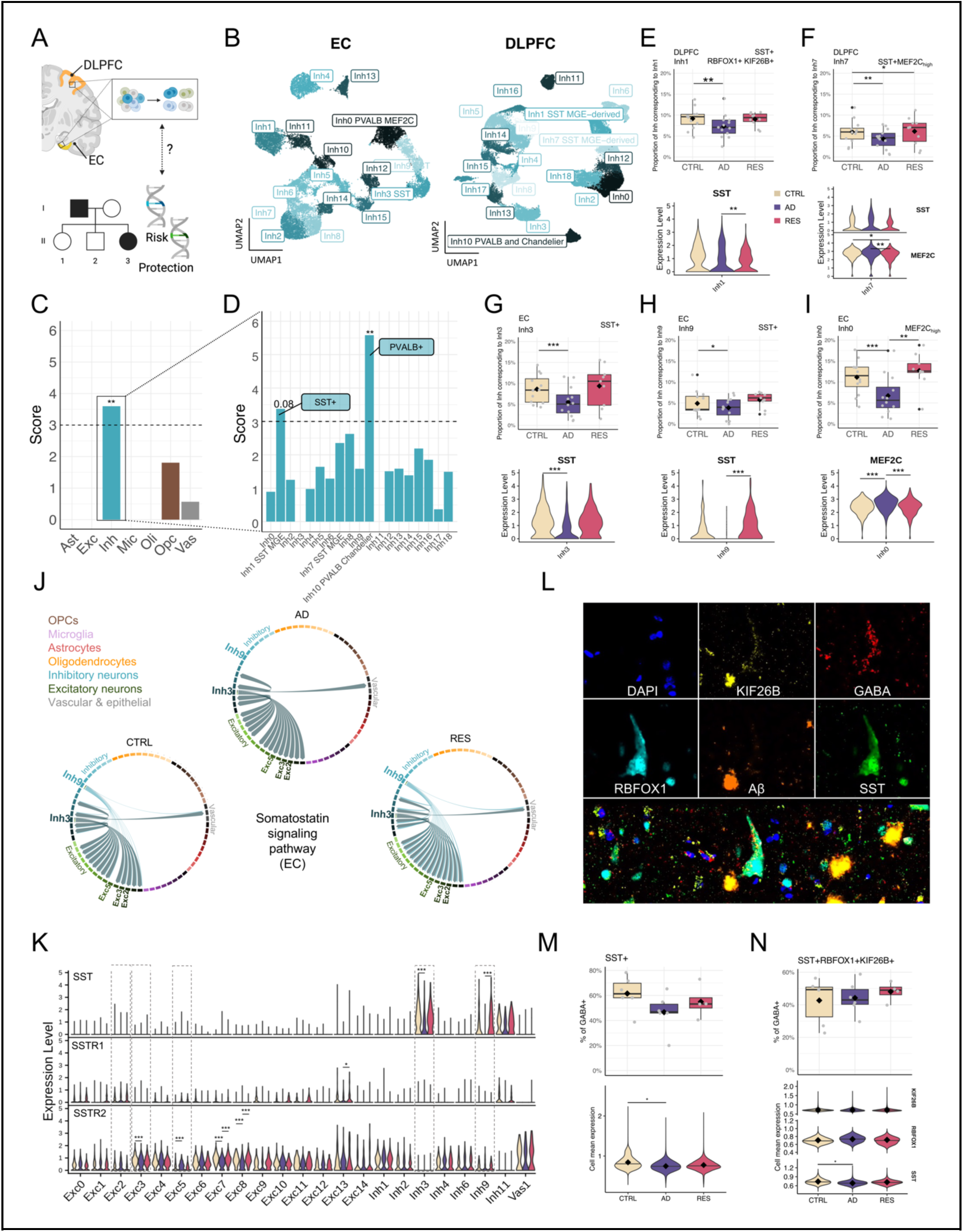
Inhibitory neurons as key players in protection against AD. **(A)** Brain regions analyzed; created with BioRender.com. **(B)** UMAP embedding of inhibitory neurons from the EC (left) and DLPFC (left). **(C-D)** Cellular enrichment results of genes annotated with protective rare variants in (C) major cell types and (D) subtypes of inhibitory neurons from the DLPFC. Y-axis shows standard deviation from the bootstrapped mean. Stars denote Bonferroni-adjusted P-values. **(E-I)** Distributions of cell proportion (top) and gene expression levels of marker genes (bottom) for the DLPFC (E) Inh1, (F) Inh7, and the EC (G) Inh3, (I) Inh9, and (J) Inh0 subpopulations. Stars show FDR-adjusted P-values from a Dirichlet multinomial regression model (**Table S18**). **(J)** Chord diagrams displaying the SST signaling pathway in cell subpopulations from the EC with significant changes per diagnostic group (**Figure S12** and **Table S17**). **(K)** Expression levels of the ligand and receptors involved in the SST signaling pathway shown in (J). **(L)** Representative image from IF staining of markers in a resilient DLPFC brain section. **(M)** Proportions of all SST+ (GABA+) cells in each subject across diagnostic groups (top). Distribution of mean SST normalized intensities in SST+ cells (bottom). N_subjects_ = 16 (6 CTRL, 6 AD, 4 RES), N_cells_ = 1,279,938 (CTRL = 434,270, AD = 559,681, RES = 285,987). **(N)** Distribution of proportions of all SST+ RBFOX1+ KIF26B+ (GABA+) cells (top). Distribution of mean intensities of each marker in SST+ RBFOX1+ KIF26B+ cells (bottom). N_subjects_ = 16 (6 CTRL, 6 AD, 4 RES), N_cells_ = 721,126 (CTRL = 261,938, AD = 315,542, RES = 143,646). Stars in top **(M)** and **(N)** indicate FDR-adjusted P-values from a Dirichlet multinomial regression. Stars in bottom (**M**) and (**N**) indicate nominal P values of a Wilcoxon test on the subject-level means; diamonds show the grand mean. *Adj.P < 0.05, ** Adj.P < 0.01, *** Adj.P < 0.001

Two DLPFC subpopulations of inhibitory neurons were enriched for genes annotated from protective rare variants (**Figure 3D**), “DLPFC:Inh1” (adj P = 0.08) and “DLPFC:Inh10” (adj-P = 0.007). Based on the analysis of subtype-specific ‘marker genes’ (see **Methods**), we characterized DLPFC:Inh1 as somatostatin (*SST)*-expression inhibitory neuron (which encodes for somatostatin) while DLPFC:Inh10 corresponded to parvalbumin (*PVALB*)-expressing inhibitory neuron. The inhibitory neuronal subpopulation “EC:Inh3” also showed enrichment for genes from protective rare variants in the EC (**Figure S10F**), and corresponds to SST+ interneurons. The expression of genes from protective variants was also enriched in several subtypes from the independent validation dataset [25], “In1” (SST+), “In2”, “In4” (SST+), and “In6”. “In9” (PVALB+) failed to pass the statistically significance threshold (adj-P = 0.11) in this additional dataset.

### Differential proportions of protection-associated inhibitory neurons

To resolve an association with resilience or resistance, we investigated changes in cell proportion for each cell subclass (Dirichlet test, **Table S15**) and compared the results with cell types enriched in rare variants. In the PVALB+ DLPFC:Inh10 subtype, the top significant change in cellular distributions, we observed a decrease in the number of these interneurons in the resilient group compared to controls (**Figure S10H**, adj-P = 0.02), suggesting that PVALB+ inhibitory neurons may be important for a distinct process of protection against AD, such as resistance to AD pathology. The SST+ DLPFC:Inh1 and SST+ DLPFC:Inh7 subpopulations, which showed enrichment for genes from ‘protective’ and ‘risk’ genetic variants, respectively (**Figure 3D** and **S9B**), also showed a change in cell proportions in AD (**Figure 3E-F**). This observation was replicated in SST+ EC:Inh3 and EC:Inh9 (**Figure 3G-H**). In our DE analyses, we found lower levels of *SST* in inhibitory neurons from the EC (**Table S14**) in ADvsRES (4th DEG ranked by p-value, log2FC = −0.52, adj-P = 3.54 x 10^-29^) and ADvsCTRL (log2FC = −0.41, adj-P = 1.79 x 10^-22^). The expression of *SST* showed significant downregulation in AD in EC:Inh3 (**Figure 3G** and **Table S18**) and EC:Inh9 (**Figure 3H** and **Table S18**), but was up-regulated in AD compared to resilient subjects (ADvsRES) in DLPFC:Inh1 (**Figure 3E** and **Table S18**). Curiously, in the HC, multiple subpopulations of both inhibitory and excitatory neurons showed an increase in the expression of *SST* in AD compared to resilient (ADvsRES, **Table S18**).

### Disrupted cell-communications of protection-associated inhibitory neurons

To further investigate how transcriptomic dynamics affect cellular dynamics in resilience, we analyzed cell-cell communication events, inferred from ligand-receptor (L-R) co-expression, using the CellChat algorithm. The number of significant inferred communication events was increased in resilient subjects in cell major populations and subpopulations in the DLPFC and EC compared to both AD and controls in most cases (**Figure S13A**, **Figure S13B**). The disruption of cell-cell communication events was marked by multiple differentially regulated pathways in ADvsRES, ADvsCTRL, and RESvsCTRL (**Figure S14)**. The somatostatin signaling pathway was predicted as changing in all brain regions in all three comparisons (**Figure S14**). In the EC, there was a complete loss of communication originating from the SST+ EC:Inh9 subtype in AD (**Figure 3J**), which could be explained at least in part by changes in the levels of *SST* in this subpopulation (**Figure 3K**). In addition to SSTR2, predicted to interact in the somatostatin pathway in all diagnostic groups, SSTR1 was identified as a receptor involved in interactions between subclasses of inhibitory and excitatory neurons from the EC solely in resilient individuals (**Table S17**). Both classes of SST+ inhibitory neurons were predicted to interact with multiple classes of excitatory neurons, including EC:Exc2, EC:Exc3, and EC:Exc5 (**Figure 3J** and **Figure 4**). Collectively, these observations support recent reports by others that SST+ inhibitory neurons are particularly vulnerable in AD, and suggest that SST dynamics may play a role in cognitive resilience.

**Figure 4.**
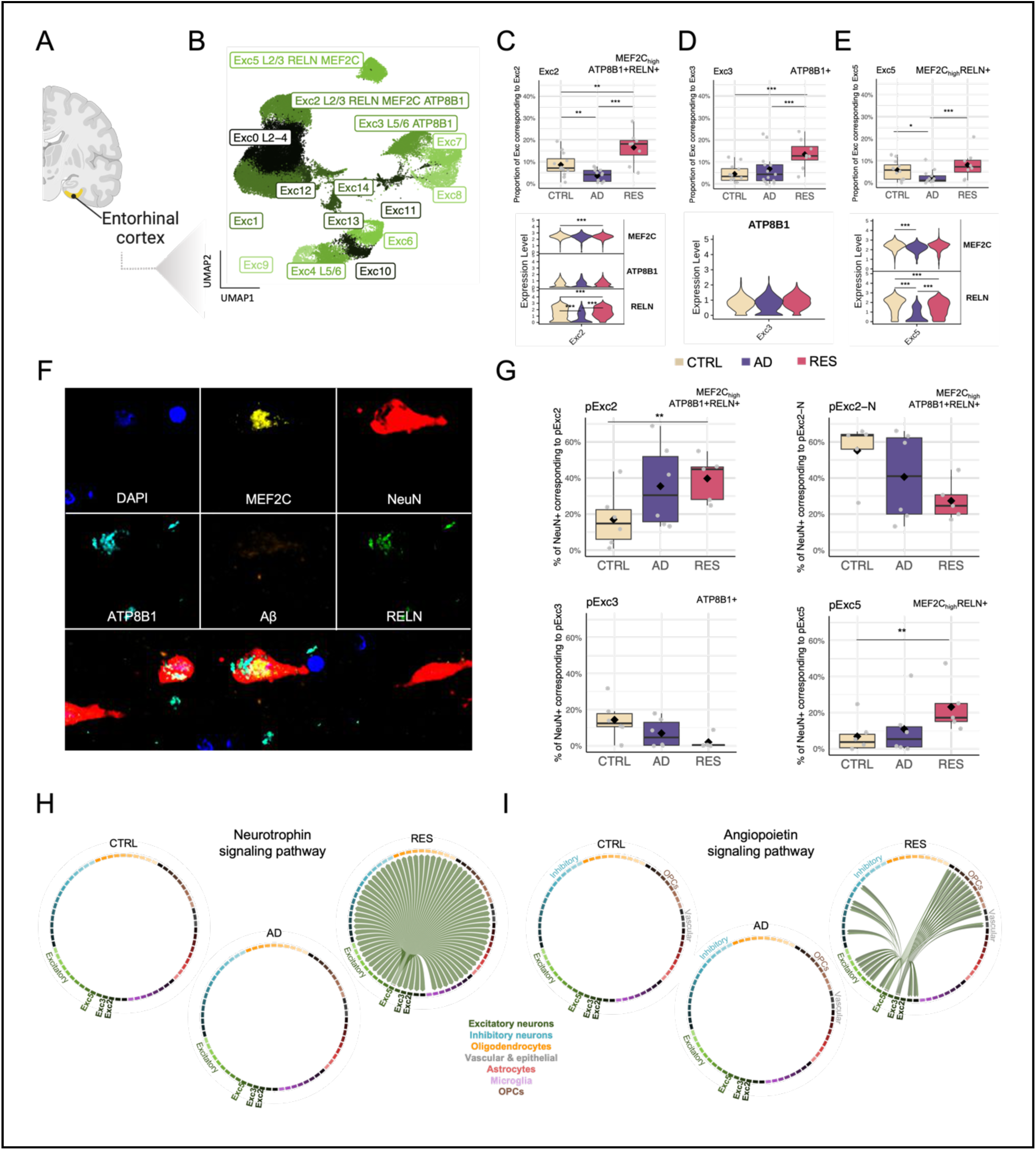
Excitatory neuronal subpopulations expressing MEF2C and ATP8B1 exhibiting resilient behavior. **(A)** Brain region for which results are shown in the figure. Figure created in part with BioRender.com. **(B)** UMAP plot showing the subclusters (‘subpopulations’) investigated in excitatory neurons from the EC, identified using the Harmony algorithm, in the ROSMAP cohort. See **Table S14** for detailed annotations. **(C-E)** Cell proportion distributions for the MEF2C_high_ ATP8B1+ RELN+ EC:Exc2 subpopulation **(C)**, ATP8B1+ EC:Exc3 subpopulation (D), and MEF2C RELN+ EC:Exc5 subpopulation **(E)**. A Dirichlet multinomial regression model was used to identify differences in cell proportions among the three diagnostic groups. P-values were adjusted using FDR correction. **(F)** Immunofluorescence representative pictures showing NeuN, RELN, MEF2C, ATP8B1, and Aβ in EC brain sections from an independent cohort. **(G)** Box plots showing cell proportion distributions for MEF2C^high^ATP8B1+ RELN+ (top; top left: positivity in the cytoplasm and cell membrane; top right: positivity in the nucleus), ATP8B1+ (bottom left), and MEF2C^high^ RELN+ (bottom right) neurons (NeuN+), identified by immunostaining. Stars indicate significance level based on FDR-adjusted P-values from a Dirichlet multinomial regression model. N_subjects_ = 17 (6 CTRL, 6 AD, 5 RES), N_cells_ = 81549 (CTRL = 19596, AD = 28107, RES = 33846) cells. **(H-I)** Chord diagrams displaying the neurotrophin (NT) signaling pathway (H) and angiopoietin (ANGPT) signaling pathway in cell subpopulations in the EC, predicted as significantly changing (Figure S12 and Table S17) from a cell-cell communication analysis based on ligand-receptor interactions. * Adj-P < 0.05, ** Adj-P < 0.01, *** Adj-P < 0.001.

### Novel candidate markers of resistance and protection-associated inhibitory neurons

Some protective rare variant-associated genes were also remarkably identified as marker genes for the SST+ DLPFC:Inh1 population. To confirm protein co-expression of these marker genes, we performed immunofluorescence imaging and mapped SST+ inhibitory neurons in human DLPFC sections from an independent cohort of formalin-fixed paraffin-embedded (FFPE) brains (see **Methods**), by staining for SST, GABA, RBFOX1, KIF26B, and Aβ (**Figure 3L**) (**Table S16**). *RBFOX1* (RNA binding fox-1 homolog 1) and *KIF26B* (kinesin family member 26B) were protective rare-variant associated gene markers which were selected based on high neuronal expression (*Human Protein Atlas)* and availability of suitable antibodies for targeting.

We identified and annotated cellular populations based upon clustering of the single-cell multiplex immunofluorescence (mIF) (see **Methods**) (**Figure S11A-C**) [28,29]. Pearson correlation analysis confirmed cellular co-expression of defining markers, establishing the potential value of rare-variant associated genes as co-factors of somatostatin expression (**Figure S11D**). The overall proportion of SST+ cells varied between AD, resilient, and control groups. Despite showing the expected decrease in AD (**Figure 3M**), a Dirichlet test failed to reach statistical significance, which may be explained by the small effect size and the low number of donors tested in this cohort (n = 4-6 per group). Nonetheless, we did observe a significant decrease in the cell-mean intensity of SST in SST+ neurons from AD and RES subjects as compared to controls (**Figure 3M**). A subpopulation of SST+ neurons, which expressed intracellular amyloid-beta (AB+) showed a significant increase in proportion in AD versus CTRL (**Figure S11F**). No differences in cell proportions were detected for a specific subpopulation that expressed all selected markers (SST+ RBFOX1+ KIF26B+ GABA+ cluster) at the protein level (**Figure 3N**).

### Region-specific resilient phenotypes in excitatory neurons

We investigated the cellular distributions of DEGs from ADvsRES in DLPFC snRNAseq data from ROSMAP [18] (expression-weighted cellular enrichment, see **Methods**) and observed that genes whose expression was decreased in AD compared to resilient (ADvsRES) were enriched in excitatory neurons (**Figure S3A-B**). In contrast, DEGs for which gene expression was increased in AD were enriched in vascular/epithelial cells (**Figure S3A-B**).

When investigating their cellular distribution, we noticed that genes whose expression was down-regulated with cognitive loss were enriched in excitatory neurons, similarly to DEGs from ADvsRES, while up-regulated genes were enriched in astrocytes (**Figure S3C-D**). Of note, two subtypes of excitatory neurons from cortical layers L2-4 (DLPFC:Exc0 and DLPFC:Exc1) enriched for down-regulated genes in ADvsRES and down-regulated genes associated with cognitive decline (**Figure S3**). This suggests molecular changes happening in these excitatory neuronal subtypes that may drive the cognitive impairment observed in AD.

Analysis of excitatory neurons identified multiple subpopulations expanded in resilient subjects in the EC (**Figure 4**), one of the first brain regions to be affected in AD. Two of these expressed *ATP8B1* as a ‘marker gene’, and two expressed *MEF2C*. These *MEF2C*_high_ populations also had a high expression of RELN (Reelin). Interestingly, MEF2C, ATP8B1 and RELN have all been previously associated with cognitive resilience [4,17,18,30,31]. One of the subtypes, EC:Exc2, annotated to cortical layers II-III, expressed both *ATP8B1* and *MEF2C* as markers. *ATP8B1* was significantly up-regulated in excitatory neurons from the EC in resilient subjects compared to controls (RESvsCTRL, **Table S13**), which we did not observe in cell subtypes (**Figure S12A**).

*MEF2C*, on the other hand, showed no significant changes in the EC in major cell types but was significantly up-regulated in AD and down-regulated in resilience in multiple neuronal subtypes from the EC (**Table S18**). *MEF2C* was also significantly up-regulated in AD and down-regulated in resilience in hippocampal neurons in major cell types (**Table S13**) and subtypes (**Table S18**).

In addition, we observed a depletion of two populations of excitatory neurons expressing CDH9+ (EC:Exc0) and RBFOX1 (EC:Exc4), respectively (**Figure S12 B**). Both subtypes of excitatory neurons were recently reported to be selectively vulnerable in AD [32]. The biggest subpopulation of inhibitory neurons in the EC, EC:Inh0 (**Figure 3I**), also exhibited decreased proportions in AD subjects. The same pattern was observed in a subcluster of inhibitory neurons in the DLPFC:Inh7 (**Figure 3F**). Interestingly, both populations of inhibitory neurons express *MEF2C* as a ‘marker gene’.

### Novel candidate markers of resilience-associated excitatory neurons

To further explore excitatory neuronal subpopulations with a potential role in cognitive resilience, we mapped MEF2C, ATB8B1, and RELN in EC sections from the independent cohort of FFPE human brains mentioned above (see **Methods**) by staining for MEF2C, ATP8B1, RELN, NeuN, and Aβ (**Figure 4F**). We followed the same procedure as in the DLPFC to preprocess the resulting mIF data, and identify and validate the presence of EC:Exc2, EC:Exc3 and EC:Ex5 populations (**Figure S12C-F**). As a quality control, we also reproduced the increase in AB+ cells in AD and RES subjects compared to CTRL (**Figure S12G**). Changes in proportions of ATP8B1+ (EC:pExc3) cells could not be replicated with immunostaining (**Figure 4G**). This discrepancy may be due to gene expression changes not always being reflected at the protein level, as well as inherent differences between RNAseq and immunofluorescence, including variations in sensitivity and detection thresholds.

When examining the proportion of NeuN+ cells corresponding to the other two subpopulations of excitatory neurons, we reproduced the expansion of MEF2C_high_ RELN+ (EC:pExc5) and MEF2C_high_ ATP8B1+ RELN+ (EC:pExc2) neurons (NeuN+) in resilient subjects, previously observed in the snRNAseq analysis (**Figure 4G**). To our surprise, we found two distinct subpopulations expressing MEF2C, ATP8B1, and RELN at the protein level (both equivalent to the EC:Exc2): pExc2 expressing similar levels of MEF2C in all cellular compartments, and pExc2-N which expressed MEF2C mainly in the nucleus (**Figure S12H**). Of the two, only pExc2 constituted a higher proportion of neurons in resilient subjects (**Figure 4G**, pExc2); whereas pExc-N showed an opposite pattern (**Figure 4G**, pExc2-N). Moreover, the levels of nuclear MEF2C among pExc-N cells were significantly higher in the RES group as compared to CTRL **(Figure S12I**). These results point to a potential role of post-transcriptional or post-translational mechanisms regulating the subcellular localization of MEF2C in resilience.

### Disrupted cell-communications of resilience-associated excitatory neurons

Inspection of cell-cell communication of the resilience-associated excitatory subpopulations revealed signaling pathways that likely play an important role in cognitive resilience (**Figure 4H-I** and **Figure S14**). Two pathways were unique to resilient subjects (i.e., were absent in control and AD): neurotrophin (**Figure 4H**) and angiopoietin (**Figure 4I**). For the neurotrophin signaling pathway, EC:Exc5 was labeled as the neuronal subtype of origin (or source), BDNF as the ligand, and NTRK2, SORT1 (sortilin 1), and NGFR (nerve growth factor receptor) as receptors. BDNF/NTRK2, critical for neuronal functions such as survival, morphogenesis, and plasticity, triggers MAPK/ERK (mitogen-activated protein kinase and extracellular signal-regulated protein kinase), PI3K (phosphoinositide 3-kinase), and PLCγ (phospholipase Cγ) [33]. *LINGO1*, the negative regulator of BDNF/NTRK2 described in the previous section as being broadly down-regulated in resilient subjects, had its expression significantly increased in AD in ADvsRES (log2FC = 0.60, adj-P = 1.26×10^-15^) and ADvsCTRL (log2FC = 0.65, adj-P = 1.23×10^-18^) in EC:Exc5, with no significant changes observed in RESvsCTRL. The angiopoietin signaling pathway (**Figure 4I**) was predicted as having EC:Exc3 and EC:Exc5 as the source cellular subpopulations, ANGPT2 (angiopoietin-2) as the ligand, and as receptors TEK receptor tyrosine kinase and ITGA5:ITGB1 (integrin alpha-5/beta-1). ANGPT2 regulates angiogenesis through TEK and integrin signaling [34]. It has been reported as neuroprotective in a model of ischemic stroke [35], suggesting that the communication between EC:Exc3/Exc5 and the subpopulation of fibroblasts (EC:Fib) is potentially related to the promotion of angiogenesis.

Additional signaling pathways showing changes in resilience (**Figure S14** and **S15**) with EC:Exc5 as source included ncWnt (non-canonical wingless-related integration site, targets: oligodendrocytes), periostin (target: EC:Inh1), BMP (bone morphogenetic protein, targets: subtypes from all major cell types), and TGF-β (transforming growth factor beta, target: EC:Fib). Of note, BMPs belong to the TGF-β superfamily, binding various TGF-β receptors, and are characterized by a mutually inhibitory crosstalk with canonical Wnt signaling [36]. Also, we observed a loss of TULP (tubby-like proteins) signaling between all three resilient excitatory neuronal subtypes (EC:Exc2, EC:Exc3, and EC:Exc5) and subtypes from multiple major cell types (**Figure S15C**). Signaling pathways observed to be lost in AD, with EC:Exc2, EC:Exc3, or EC:Exc5 as sources, are shown in **Figure S16**, and included, for example, KIT (or c-Kit) and EGF/EGFR signaling. Intriguingly, we noticed a shift of the EGF signaling pathway from EC:Exc2 as the source in controls to EC:Exc3 in resilience (**Figure S16C**, ligand: BTC; receptors: EGFR and ERBB4).

## Discussion

Using multimodal integration of molecular and cellular signatures, we have defined molecular determinants of protection and resilience against AD across brain tissues, cellular populations, and communication. Resilient and resistant brains protect cognition through a combination of synaptic plasticity, selective survival of SST+ inhibitory neurons, and increase in their excitatory neuron populations. They also upregulate protein homeostasis, reduce neuroinflammation, and activate astrocytic responses to AD pathology.

At the tissue level, transcriptomic changes occur at onset of AD dementia, but resilient individuals show minimal differences between age-matched healthy controls and presymptomatic individuals. Resilient individuals maintain neuronal diversity, synaptic markers, and cortical thickness similar to healthy, pathology-free controls, suggesting resilience reflects an intermediate phase of AD progression.

Tissue-level transcriptomic differences between resilience and healthy individuals are marked by upregulation of *GFAP* and downregulation of *KLF4*. GFAP, key marker for reactive astrocytes [37], increases in expression with AD progression and may reflect early astrocytic activation in resilient brains, supported here by snRNAseq analysis. Investigation of cognitive decline in AD should account for GFAP activity in assessing mechanisms of resilience.

Resilience-dependent loss of *KLF4* expression, a nuclear transcription factor in microglia and endothelial cells, appears to be restored in AD individuals. Lipopolysaccharide (LPS) stimulation increases *KLF4* expression in microglia, while its knockdown reduces pro-inflammatory cytokines [38]. KLF4 expression also rises in microglia exposed to oligomeric Aβ42 [39] and its downregulation promotes axonal regeneration after optic nerve injury [40]. Additionally, KLF4 targets are dysregulated in AD and linked to anti-inflammatory roles in brain endothelial cells. These findings suggest that KLF4 is involved in multiple resilience-associated regulatory processes in AD.

At cellular resolution, we have identified pervasive reorganization of protein folding and degradation processes associated with cognitive resilience across distinct cellular populations (**Figure 2**). Resilience-related re-organization is characterized by widespread down-regulation of Hsp90, contrasted by selective up-regulation of Hsp40, Hsp70, and Hsp110 in excitatory neurons, exhibiting patterns opposite to those seen in AD. Hsp90, a key regulator of protein folding and degradation, binds tau at aggregation-prone repeats, mediating its stabilization and degradation [41]. Hsp40 h counteracts formation and propagation of toxic tau aggregates [42]. Hsp70 has been shown to inhibit tau and sequester its aggregates with high affinity while seemingly contrasting reports point to its AD upregulation in the superior frontal gyrus. [43,44]. *Hsp110* knockout mice have been reported to exhibit pTau accumulation and neurodegeneration [45]. Differential regulation of molecular chaperones suggests that cognitive resilience involves distinct rearrangement of protein folding and degradation dynamics.

Genetic clues from common AD-associated variants, including expression of their corresponding genes, have pointed to a major role for microglia and immune processes in AD [46]. We have shown that common and rare AD-associated mutations may impact different populations of cells [23]. Rare variants differ from common, environmentally influenced variants as they may be more likely to have a direct effect on molecular processes [47,48]. We show that genes associated with rare ‘protective’ and ‘risk’ genomic variants are significantly biased in their expression towards subtypes of human cortical neuronal cell populations;SST+ and PVALB+ inhibitory neurons (**Figure 3D**). SST+ neurons show vulnerability in AD (**Figure 3E-G**). SST+ and PVALB+ interneurons regulate the activity of excitatory pyramidal neurons. Both have been associated with AD [15,16,49]. In APP/PS1 mice, SST+ interneurons are hyperactive, while PVALB+ interneurons are hypoactive [50]. Rare genomic variants may illustrate a resistance phenotype (rather than resilience) in inhibitory neurons, highlighting potential mechanisms that drive protection and risk prior to manifestation of advanced AD pathology.

AD-associated rare variants were differentially enriched in two key populations of SST+ interneurons derived from the medial ganglionic eminence (MGE) in the DLPFC. DLPFC:Inh1 showed enrichment for genes associated with protective variants, while DLPFC:Inh7 was enriched for risk variants (**Figures 3D** and **S9**). Both cellular populations displayed significant changes in proportions in AD. Protective rare variant enrichment was replicated in an SST+ population of interneurons in the EC, EC:Inh3. Another SST+ subpopulation, EC:Inh9, also displayed significant changes in cell proportions in AD. Additionally, a PVALB+ subpopulation of cortical inhibitory neurons, DLPFC:Inh10 showed the most significant cellular enrichment of genes from protective AD-associated rare variants. This population showed a decrease in the proportion of cells in resilience compared to controls, suggesting a role for PVALB+ neurons in resistance and protection against AD.

We identified genes associated with rare genomic variants that define resistance-linked inhibitory neuronal populations. *RBFOX* and *KIF26B* appeared as potential markers of the protection-associated SST+ DLPFC:Inh1 population of interneurons (**Figure S11**). *RBFOX1* functions as a neuronal RNA-binding protein that regulates alternative splicing and the stability of mRNAs regulating mRNA stability and synaptic transmission [51]. Influencing neuronal development and synaptic networks, it has been genetically associated with brain amyloidosis in AD [52]. Reduced expression of *RBFOX1* has been shown to occur with higher Aβ burden and cognitive decline [53]. RBFOX1 is specifically expressed in SST and Parvalbumin neurons and plays a known role in regulating the developmental integration of SST interneurons into cortical circuits, influencing connectivity [54]. Immunofluorescence image analysis confirmed significant co-expression of SST and RBFOX1 across multiple compartments of neuronal populations. Using network analysis, we confirmed that RBFOX1 interacts with several other rare variant-associated risk and protective genes that were also identified as marker genes of the DLPFC:Inh1 SST+ subpopulation, including genes enriched in cell morphogenesis, paranodal junction, potassium channel complex, and axonogenesis (**Figure S17** and **Tables S19-20**). Our results suggest that RBFOX1 expression is preserved in resilient individuals in SST+ interneuron populations.

KIF26B, co-expressed with RBFOX1, is an intracellular motor protein, is thought to participate in cell signaling. *KIF26B* is a member of a gene co-expression module showing early downregulation in vulnerable SST+ neurons [15]. Its protein levels are reduced within AD CA1 pyramidal neurons, [55]. We noticed lower gene expression of *KIF26B* in AD compared to resilient individuals (ADvsRES) in hippocampal excitatory neurons (**Table S13**). We also observed a decrease in the expression of *KIF26B* in AD (ADvsRES and ADvsCTRL) in inhibitory neurons in the DLPFC and EC (**Table S13**).

Our study contributes to the growing body of evidence suggesting critical involvement of SST+ interneurons in the pathogenesis of AD. Cortical and hippocampal SST+ interneurons are selectively vulnerable in AD neurodegeneration [15–17,49,56], and are associated with cognitive resilience [16,17,56,57]. The genomic locus of SST has been associated with risk of developing AD [58,59]. *SST* expression declines throughout brain aging and AD pathogenesis, partially explained by aging-associated hypermethylation of its promoter [60–62]. SST+ inhibitory neurons colocalize with Aβ, where SST (usually co-released with GABA) interacts with Aβ to promote degradation. Aβ-SST interaction requires that at least one of the binding molecules is in a pre-aggregated oligomeric form [reviewed in [63]]. SST expression in resilience, possibly in a monomeric form, may exert a protective role by facilitating Aβ degradation and clearance.

A PVALB+ population of inhibitory neurons, enriched for genes linked to protective rare variants, may be associated with resistance to AD pathology as we observed no resilience-related changes in their proportion dynamics. Studies have identified PVALB+ interneurons as resistant to AD pathology [64] and neurodegeneration [65]. While recent findings report vulnerability to increased beta-amyloid and pTau [15], a separate study identified a higher proportion of PVALB+ inhibitory neurons in individuals with cognitive decline [16], similar to our findings (**Figure S10H**).

We have found an excitatory neuronal basis for the resilience phenotype. In marked contrast with the resistance phenotype in inhibitory neurons, we identified several excitatory neuronal subtypes exhibiting resilience to AD (**Figure 4**). *ATP8B1*, *MEF2C,* and *RELN* emerged as key marker genes within these resilient subpopulations (Exc 2, Exc 3, and Ex 5 in EC), with *ATP8B1* showing significant up-regulation in gene expression in resilient subjects compared to controls. This upregulation is consistent with previous findings associating *ATP8B1* and *MEF2C* with cognitive resilience in AD [4,30]. *ATP8B1* was primarily up-regulated in excitatory neurons from the EC, whereas *MEF2C* displayed significant up-regulation in AD and down-regulation in resilience across multiple neuronal subtypes from both the EC and HC. Beyond excitatory neurons, *MEF2C* was also a marker of subpopulations of inhibitory neurons in the EC and DLPFC showing decreased proportions in AD. This suggests that the role of MEF2C in cognitive resilience may extend beyond excitatory neurons, possibly contributing to the maintenance of excitatory/inhibitory balance in key brain regions. In the multi-region atlas of AD brains, *RELN+* L2/3, L2 and L3 excitatory neurons in EC were shown to be selectively vulnerable to AD and thus potentially associated with resistance. A genetic variant of *RELN* was recently shown to be associated with extreme resilience in autosomal dominant AD [31]. Our findings highlight intricate roles for these excitatory neurons expressing *ATP8B1*, *MEF2C*, and *RELN* that are both selectively vulnerable to AD but showing increased proportions in resilient subjects, suggesting involvement in both resilience and resistance (**Figure 4**). Using immunostaining in an independent cohort of FFPE human brain sections, we examined gene expression at the protein level using. RELN and ATP8B1 co-expression was confirmed in NeuN+ cells. MEF2C expression did not correlate with either RELN or ATP8B1 at the cellular level. Subcellular relocation of MEF2C occurred in neuronal populations in resilience.

Neuroplasticity in the DLPFC, and consequently working memory performance, is impaired in AD [66]. In AD dementia, cortical hyper-excitability is inversely correlated with overall cognition and executive functions [67]. The degeneration of SST+ interneurons likely disrupts the excitatory/inhibitory balance by impairing the inhibitory modulation of pyramidal neurons. This loss of inhibitory neuron function destabilizes neuronal networks, leading to hyperexcitability [68] and contributing to the cognitive deficits observed in AD. In contrast to inhibitory neurons, changes in cell proportions for excitatory neurons were only detected in the EC. We propose that in the DLPFC and HC, inhibitory neurons exhibit selective vulnerability to AD, while excitatory neuron loss may occur more broadly.

Cell-cell communication analysis highlighted neuronal cell interactions in resilience particularly through neurotrophin and angiopoietin signaling pathways. BDNF and its receptor, TrkB (NTRK2) (neurotrophin pathway) are essential for neuronal survival, synaptic plasticity, and neurogenesis [33]. BDNF/NTRK2 coexpression was uniquely active only in resilient subjects, suggesting a protective role in AD. NTRK2 and its interaction partner NGFR, are linked with protective rare variants. BDNF supports hippocampal neurogenesis by differentiation and survival of new neurons, processes disrupted in AD. *LINGO1*, a negative regulator of BDNF/NTRK2, was up-regulated in AD but not in resilient individuals. LINGO1 promotes APP degradation, contributing to Aβ deposition Aβ [69,70] and its up-regulation is implicated in AD and Parkinson’s disease [71,72]. Anti-LINGO-1 antibodies improve cognition, neurogenesis, and synaptic protection in mice [73], and increase the numbers of GABAergic interneurons [74]. This study is the first to report the association of LINGO1 with cognitive resilience in humans. Inhibiting LINGO1 may support neuronal function and cognitive resilience.

The angiopoietin signaling pathway, ANGPT2, TEK, and integrins ITGA5:ITGB1 were uniquely active in resilient subjects. ANGPT2 supports neuroprotection in ischemic stroke models [35]. Angiopoietin signaling between excitatory subpopulations and fibroblasts suggests that angiogenesis could maintain cognitive function in AD pathology. We have identified ANGPT2/TEK signaling originating from excitatory neurons targeting excitatory and inhibitory neurons and oligodendrocyte precursor cells (OPCs) suggesting a neuroprotective role in AD.

To encapsulate the molecular and cellular events we have discovered here, we have developed a model of cellular processes and pathways driving cognitive resilience (**Figure 5**). In the resilient response to cellular stress by AD related pathology, LINGO1 is downregulated [72,75,76]. LINGO1 inhibits NTRK2/TrkB through phosphorylation and so inhibits BDNF binding to NTRK2/TrkB. Downregulation of LINGO1 releases BDNF inhibition [77,78] which leads to autophosphorylation of the NTRK2/TrkB receptor [79], activating downstream signaling cascades, including ERK5 [80] which, in the nucleus, activates the transcriptional activity of MEF2C and MEF2 through phosphorylation (**Figure S18**). MEF2C, which in our study locates in the nucleus when in resilient cells, shows transcriptional levels positively associated with cognitive ability. Mef2 target genes are significantly overrepresented among the genes that are most predictive of cognition. Mef2c may play a significant role in cognitive resilience against AD by regulating genes whose expression is critical for neuronal survival, synaptic plasticity, and reduction of hyperexcitability [12]. MEF2C has been shown to modulate KLF4 [81]. KLF4 acts as a transcription factor involved in neuroprotection, particularly through anti-apoptotic pathways [82]. ERK5/KLF4 signaling is a common mediator of neuroprotective effects seen in neurons exposed to nerve growth factor (NGF) and oxidative stress [82]. Importantly, BDNF/TrkB interaction affects synaptic plasticity but also modulates pathways tied to neuronal excitability, which can be disrupted by tau hyperphosphorylation, a process known to decrease BDNF expression [83,84].

**Figure 5.**
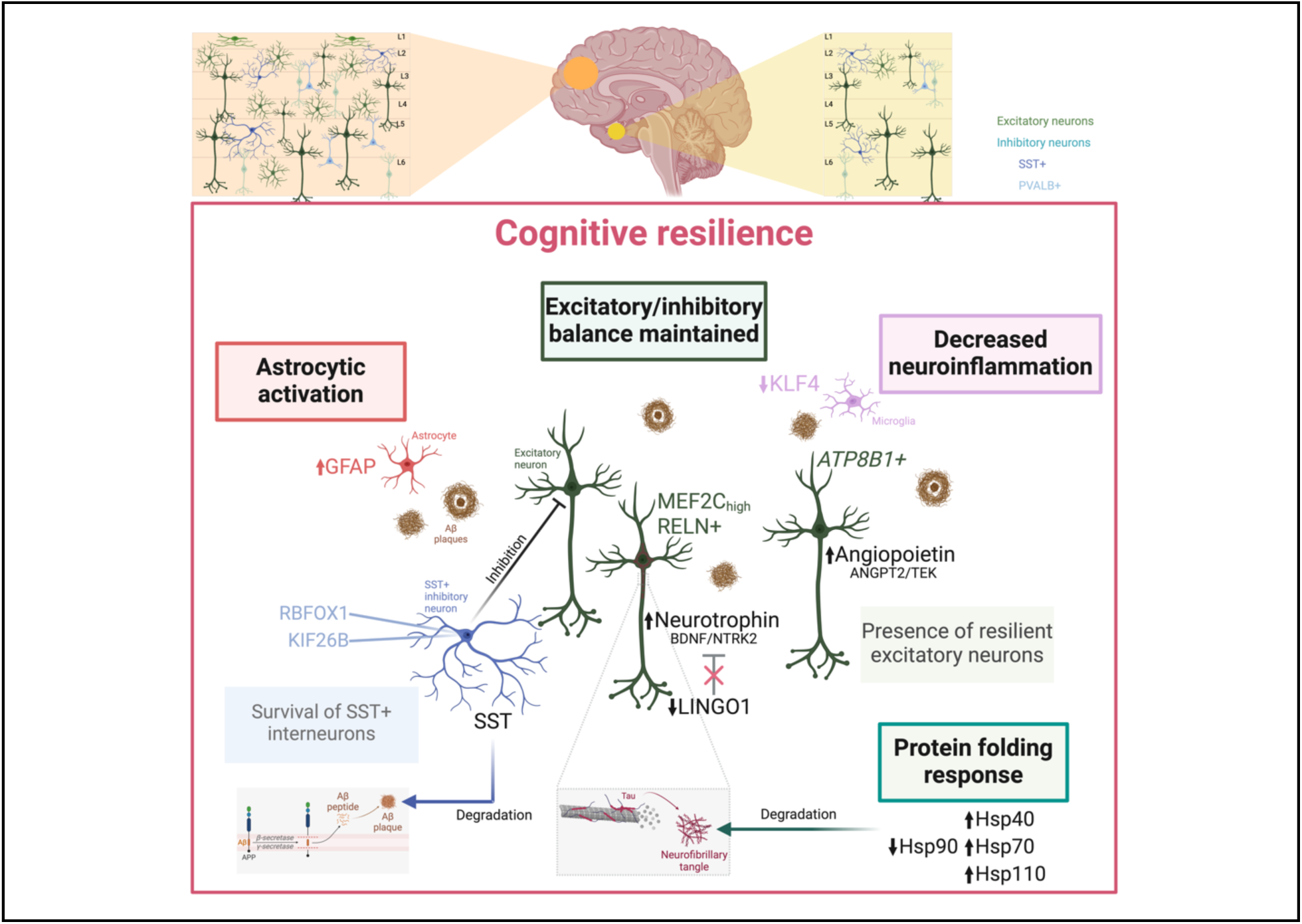
A functional model of resilience. Our model proposes that cognitive resilience is driven by the maintenance of the excitatory/inhibitory neuronal balance (dark green), sustained by resilient excitatory neurons expressing MEF2C and ATP8B1. These neurons engage in resilience-relevant signaling pathways, including neurotrophin (BDNF/NTRK2), modulated by the down-regulation of LINGO1, and angiopoietin (ANGPT2/TEK). Protein folding and degradation processes are reorganized in resilience, with increased expression of Hsp40, Hsp70, and Hsp110 in excitatory neurons and down-regulation of Hsp90, enhancing the degradation of pathological tau (mint green). SST+ inhibitory neurons, typically vulnerable in AD, are preserved in resilience, including subpopulations expressing RBFOX1 and KIF26B (blue), contributing to the balance of neuronal excitation. Additionally, SST release from these neurons promotes the degradation and clearance of pathological Aβ. In terms of glial response, resilience shows astrogliosis marked by increased GFAP in astrocytes (red), a feature shared with AD. However, it contrasts with AD by exhibiting a reduction or absence of microglial activation, characterized by decreased KLF4 expression, leading to reduced neuroinflammation (pink). Figure created with BioRender.com.

### Limitations

Definitions of the (molecular) resilience phenotype in AD vary signficantly across studies [17,18,85–88]. In this study, we adopted a discrete but extreme definition of resilience, selecting individuals with advanced amyloid and tau pathology at death but preserved cognitive function. To stratify individuals who may eventually develop dementia if they lived long enough, we could have analyzed continuous longitudinal changes in cognition. Without this approach, we were unable to capture resilience as deviations from expected cognitive trajectories, including moderate levels of resilience. We have not considered how other pathologies and their readouts, such as neuroinflammatory markers, contribute to dementia and resilience Other limitations of our study include the inherent challenges of working with rare genetic variants, which were identified through a familial genetic analysis and whole genome sequencing. Due to their rarity, these variants have not achieved genome-wide significance, which limits their statistical power. This limitation was partially addressed in our initial study by validation in an independent case-control whole genome sequencing (WGS) dataset [23]. Nonetheless, we have shown that using a rank-based approach for selecting the most significant variants, followed by systems analysis, yields informative biological functions and cell type-biased expression patterns.

### Future directions

To fully define and understand resilience and resistance to AD, future studies should map cellular identities to their precise spatial locations and integrate molecular determinants of resilience at the sites of activity. Study should identify the specific cell types where resilience processes are active, their locations, and their communication networks. By understanding these relationships, we can pinpoint which cells and mechanisms should be targeted for protection to halt disease progression. Equally important, is exploring these processes at the subcellular level, particularly within synapses, where early degenerative changes often occur. The role of synaptic health in resilience is critical, as synaptic dysfunction is one of the earliest hallmarks of AD [89]. Future work should additionally investigate how sex differences impact resilience to AD, as sex-specific pathways may play a role in neuroinflammatory regulation, cognitive decline, and overall disease progression. Addressing these dimensions will offer a more complete picture of resilience mechanisms and provide new insights for therapeutic interventions.

### Conclusions

This study advances our understanding of cognitive resilience by revealing specific molecular and cellular processes that may protect against AD. Our study has identified molecular and cellular hallmarks of cognitive resilience in Alzheimer’s disease (AD), emphasizing the preservation of excitatory/inhibitory balance as critical to resilience.

Through integrative transcriptomic and single-cell analyses, we have uncovered global reorganization of protein folding and degradation pathways, highlighted by selective upregulation of Hsp40, Hsp70, and Hsp110 families and downregulation of Hsp90 in excitatory neurons. Key excitatory neuronal subpopulations, including MEF2C-high neurons in the entorhinal cortex, demonstrated resilience-associated neurotrophin and angiopoietin signaling mediated by the BDNF/NTRK2 and ANGPT2/TEK pathways, implicating LINGO1 downregulation as a novel mechanism promoting cognitive preservation.

The identification of resilient subpopulations of excitatory and inhibitory neurons, as well as key glial cell responses, highlights specific cellular mechanisms that can be leveraged for therapeutic intervention. This study exposes potential novel strategies that emulate natural resilience processes to protect against AD dementia and offers new avenues for the prevention of neurodegeneration and cognitive decline.

## Methods

### Human Subjects

Clinical, pathologic and omic data are from participants in the Religious Orders Study or Rush Memory and Aging Project (ROSMAP). All ROSMAP participants enrolled without known dementia and agreed to detailed clinical evaluation and brain donation at death [90]. Both studies were approved by an Institutional Review Board of Rush University Medical Center. Each participant signed an informed consent, Anatomic Gift Act, and an RADC Repository consent. The evaluation includes 21 cognitive performance tests, 17 of which are summarized as global cognition and five cognitive domains [91]. The tests also inform on clinical diagnoses of AD dementia and mild cognitive impairment (MCI), and the reference no cognitive impairment (NCI) [92–94]. The neuropathologic assessment includes pathologic AD based on CERAD neocortical neuritic plaque estimates the severity and distribution of neurofibrillary tangles by Braak Stage; there are also quantitative measures of amyloid-β load and PHFtau tangle density derived by 8 brain regions [95–97]. Other neurodegenerative and vascular diseases are also documented [98,99]. The brain omics data used in this study are described in detail below.

### Bulk RNAseq data analysis

#### Data retrieval and normalization

RNAseq profiles (n = 631) of dorsolateral prefrontal cortex (DLPFC) tissues of individuals from ROSMAP participants were accessed through the AMP-AD knowledge portal (syn2580853) [19,100]. Read counts of samples with non-missing covariates were analyzed. The analysis was limited to genes with non-missing length and GC content, with at least one count per million read across all the samples. Samples were normalized for library size, GC content of genes, and genes’ lengths using the *Conditional Quantile Normalization* (CQN) method [101].

An iterative Principal Component Analysis (PCA)-based approach was used to determine significant non-biological, demographical, and technical confounding covariates (e.g., RNAseq quality metrics determined by the AMP-AD consortium using Picard Tools). The PCA-based analysis was performed using protocols previously described in the AMP-AD cross-cohort harmonization study [100]. Covariates that significantly correlated with the principal components of the gene expression data were selected for adjustment in downstream analysis. The normalized RNAseq read counts were adjusted by determining the residual gene expression values from a linear model containing confounding covariates, implemented using the Limma-Voom method [102]. The final set of covariates for adjustment included: batch, sex, RNA Integrity number (RIN), %coding bases in each sample, %intergenic bases in each sample, post-mortem interval (PMI), age at death, and %pass-filter reads aligned.

#### Sample classification

Using levels of Aβ plaques and neurofibrillary tangles, and presence/absence of cognitive impairment, subjects with gene expression data available were classified into three major categories: AD (n = 187; diagnosis of AD dementia with no other cause of cognitive impairment; moderate/frequent plaques; Braak Stage III-VI), Resilient (RES, n = 68; “no cognitive impairment”; moderate/frequent plaques; Braak Stage III-VI; age > 80), Control (CTRL, n = 44; “no cognitive impairment”; sparse/none plaques; Braak Stage 0-II). An additional group of subjects was defined as Presymptomatic (PRE, n = 83, “mild cognitive impairment with no other cause of cognitive impairment”, moderate/frequent plaques; Braak Stage III-VI). All other subjects that did not fit the criteria described above were classified as “Other” (n = 249).

#### Differential gene expression analysis

Differentially expressed genes (DEGs) were determined using Limma-Voom using the following comparisons: ADvsRES, ADvsCTRL, RESvsCTRL, ADvsPRE, PREvsCTRL, and RESvsPRE. DEGs were determined by adjusting for the confounding covariates determined in the previous step. P-values were adjusted for multiple comparisons using the False Discovery Rate (FDR) method [103]. Given that transcriptomic changes in DLPFC are more moderate compared to other brain regions [104], DEGs were determined using FDR < 0.1 and |log2FC| > log2(1.1) cut-offs (at least 10% difference in absolute expression).

#### Ordinal Categorical Analysis of loss of cognition

To determine genes associated with cognitive loss, we investigated an association between gene expression and loss of cognition using a Proportional Odds Model (POM). Gene expression levels were modeled as a function of cognitive status (ordinal categorical variable dependent on gene expression), Aβ pathology (plaques), and tau pathology (neurofibrillary tangles). The analysis was performed for each gene, implemented by a Proportional Odds Logistic Regression model using the VGAM library [105,106]. All ROSMAP subjects with a cognitive status of “No cognitive impairment”, “Mild cognitive impairment with no other condition contributing to CI”, and “Alzheimer’s dementia with no other condition contributing to CI” (NINCDS/ADRDA “Probable AD”) were included in the analysis. Gene expression values were adjusted for confounding covariates (described above) prior to ordinal categorical analysis. Two-sided p-values associated with gene expression were adjusted for multiple hypothesis comparison using the Benjamini-Hochberg FDR method [103]. Additional ordinal categorical analyses were performed to determine genes associated with i) Aβ pathology (CERAD score), ii) tau pathology (Braak stage), and iii) loss of cognition without accounting for pathology levels.

#### Pathway Activity Analysis

To determine functional dysregulation events associated with AD and resilience, the PanomiR package was used to define differentially regulated pathways from the ROSMAP RNAseq data [21]. PanomiR uses a non-parametric rank-based summarization technique to generate pathway activity profiles from gene expression data [21,22,107–109]. In brief, the activity of a pathway in each sample represents the average squared ranks of the genes belonging to the pathway. PanomiR uses linear models via the Limma software package to compare pathway activity profiles between disease conditions and adjust for confounding covariates. In addition, PanomiR determines co-expressed modules of differentially regulated pathways by using network clustering algorithms and the Pathway Co-expression Network (PCxN) [109].

PanomiR was applied to the normalized ROSMAP gene expression data to map the activity of 1329 canonical pathways from the Broad Institute’s Molecular Signatures Database (MSigDB, C2, V6.2, July 2018). Dysregulated pathways were determined by comparing AD and resilient subjects using the *Limma* package [110], adjusted for confounding covariates determined in previous steps. Pathway-dysregulated p-values were adjusted for multiple-hypothesis testing using Storey’s q-value method [111]. Pathways with significant dysregulation (q-value < 0.1) between AD versus resilient subjects were mapped to PCxN. The mapping of dysregulated pathways to the PCxN network was performed using the default parameters provided in the PanomiR package, i.e., the network only contained edges corresponding to a correlation of 0.316 (sqrt(.1)) with a significance threshold of FDR < 0.05, excluding nodes without any edges in the PCxN network. Clusters of dysregulated pathways were determined using the *Label Propagation* algorithm, implemented in the *igraph* package [112].

### Single nuclei RNAseq data

#### Data processing

The generation of the single nuclei RNAseq (snRNAseq) dataset covering multiple brain regions from 48 individuals from ROSMAP is described in its original publication [18,113]. Clusters corresponding to major cell types were defined across brain regions as initially reported, subcell types were re-annotated for each brain region separately (described below). Counts for protein-coding genes were extracted from snRNAseq pre-processed data (filtered for mitochondrial and ribosomal RNA, and filtered for doublets, as described in [18] and reanalyzed independently for each brain region. Quality control (QC), additional filtering, normalization, and scaling were performed using the package *Seurat* [29](version 4.1.1) in R (version 4.1.2).

#### Clustering and annotations

Brain region-, cell type-specific subclusters (unsupervised) were generated using Harmony [114] (version 1.0, resolution = 0.5), with integration by subject. Cluster annotations were generated by a combination of reference mapping and manual curation. Cells from each brain region were mapped separately to two brain reference datasets, the 10x Whole Human Brain [115], and the Human MTG SEA-AD [116], using the web tool MapMyCells [117], and compared to the annotations previously described [18]. The mapping was performed using the Hierarchical algorithm. Final annotations of each cell subtype cluster were curated manually considering the results obtained from both references and marker genes, identified as described below.

### Sample classification

Similarly to bulk RNAseq, using detailed phenotyping information from ROSMAP, subjects were classified as AD (Braak III-VI, CERAD “definite AD” or “probable AD”, and consensus cognitive diagnosis “Alzheimer’s dementia and no other cause of cognitive impairment”), RES (Braak III-VI, CERAD “definite AD” or “probable AD”, and consensus cognitive diagnosis “no cognitive impairment”), and CTRL (Braak 0-II, CERAD “no AD”, and consensus cognitive diagnosis “no cognitive impairment”). Subjects outside this criteria, classified as Other, were excluded from further analyses. We selected the dorsolateral prefrontal cortex (DLPFC) as the main focus for our initial investigations for consistency with bulk data, and we also investigated the entorhinal cortex (EC) and hippocampus (HC) as brain regions affected early in AD. Number of subjects and cells per group is shown in **Table S11**.

#### Differential gene expression analysis

Differential gene expression analyses were performed by implementing a statistical model by group using the MAST statistical framework [118](two-part generalized linear model), with a random effect for individual [119], integrated with the *Seurat* (version 4.1.1) workflow (*MAST* R package version 1.20.0). The following covariates were included in the differential expression analysis: sex, age at death, and number of unique molecular identifiers (UMIs). To find gene expression differences between the groups, the following comparisons were performed: ADvsRES, ADvsCTRL, and RESvsCTRL. All other parameters were set as default, except logfc.threshold (logfc.threshold = 0.05), with final sets of DEGs determined using |log2FC| > 0.2. P-values were adjusted based on Bonferroni correction, as per Seurat’s recommendations (satijalab.org/seurat/), using all genes available in the dataset. Genes classified as mitochondrial genes were removed from the results tables and corresponding visualizations. In parallel, marker genes for each cell subtype, i.e., genes that define each subcluster, were identified via differential expression using the function *FindAllMarkers* from the package *Seurat* (version 4.1.1) in R (version 4.1.2) [29], with the following arguments: only.pos = TRUE, min.pct = 0.25, logfc.threshold = 0.25.

#### Gene ontology enrichment analysis

Gene ontology enrichment analysis and protein-protein interaction enrichment analysis were performed using Metascape [120]. The expressed genes from each brain region-specific major cell type were used as the background list for each inquiry. The following pathway catalogs were inquired: “GO Biological Processes”, “BioCarta Gene Sets”, “Canonical Pathways”, “Reactome Gene Sets”, “KEGG Pathway”, “WikiPathways”, “PANTHER Pathway”. Protein-protein interaction enrichment analysis was carried out with the following databases: STRING, BioGrid, OmniPath, and InWeb_IM. Up and down-regulated genes were investigated separately.

#### Cellular distributions

To identify changes in cell composition between AD, Resilient, and Control, we used a Dirichlet multinomial regression model, while accounting for the proportions of all of the other cell subsets within each major cell type. Changes in cell proportions and its associated p-values were determined using the *DirichReg* function in the *DirichletReg* (version 0.7-1) R package as described by others [121]. P-values were adjusted for multiple comparisons using the Benjamini-Hochberg (FDR) correction. Subclusters with a low number of cells (counts < 50 in either of the groups) were removed from the analysis.

#### Intercellular communication

To investigate changes in intercellular communication between and within the various cell populations, we used the R package *CellChat* [122] (version 2.1.2) and followed its standard analysis pipeline with default settings.

### Whole genome sequencing data

To evaluate the cellular enrichment of genes from AD-associated rare variants, we have accessed summary statistics from a recent systematic analysis of rare variants associated with AD in two large whole-genome sequencing datasets [23]. The full analysis is described in the original publication. Briefly, single variant analysis and grouped rare variant analysis were performed in a family-based WGS dataset of 2247 subjects from 605 AD families and 1669 unrelated individuals. We have used two sets of results: a) based on single rare variants from an extended supplementary table 12 and b) based on grouped rare variants from supplementary table 15 [23]. Effect direction was identified from the Z-score for single rare variants and as the direction of the signal for the most significant single variant in the region-based analysis. Genetic variants found in asymptomatic family members were classified as ‘protective’, and genetic variants associated with AD were classified as ‘risk’ variants.

### Expression weighted cell type enrichment analysis

To investigate cellular enrichment of genes annotated from rare variants identified from WGS, we integrated snRNAseq data with rare variant-associated genes using expression-weighted cell type enrichment analysis using the R package *EWCE* (version 1.4.0) [123] with default settings, including Bonferroni correction of multiple comparisons. For computational feasibility, we partitioned the data using the R package caret (representing each subject equally), and ran EWCE in 60% of the snRNAseq data, using 10,000 permutations. Subclusters with a low number of cells (counts < 50 in either of the groups) were removed from the analysis. Genes from common variants were obtained from Bellenguez et al. 2022 (Table 1 and Table 2 from the original publication) [9].

### Gene network prediction

To determine likely interactions and functional enrichment of marker genes expressed in SST vulnerable inh1 neurons. Genes associated with unique directional variants in Table S12 were defined as follows: Rare genetic variants found in asymptomatic family members were classified as ‘protective’, and genetic variants associated with AD were classified as ‘risk’ variants (see Methods). Overlapping genes that were expressed as markers in DFPLC:inh1 in resilient subjects were selected for network generation. A network of interacting node partners was generated using the STRING protein query feature of Cytoscape 3.10.2 set at default evidence.

### Definition of risk and protective variants

The full analysis is in the original publication. Briefly, single variant analysis and grouped rare variant analysis were performed in a family-based WGS dataset of 2247 subjects from 605 AD families and 1669 unrelated individuals. We have used two sets of results: a) based on single rare variants from an extended supplementary table 12 and b) based on grouped rare variants from supplementary table 15. Effect direction was identified from the Z-score for single rare variants and as the direction of the signal for the most significant single variant in the region-based analysis. Genetic variants found in asymptomatic family members were classified as ‘protective’, and genetic variants associated with AD were classified as ‘risk’ variants.

### Immunostaining

Human brain tissue used for immunostaining experiments was collected at Beth Israel Deaconess Medical Center (BIDMC) upon autopsy. Its use was approved by the BIDMC Institutional Review Board (IRB). Clinical and neuropathological phenotypes were used to assess each individual, including cognitive status proximate to death. Brains from age- and sex-matched de-identified individuals were classified into AD (Braak V-VI, CERAD “Moderate” or “Frequent”, and a clinical diagnosis of AD dementia), Resilient (Braak V-VI, CERAD “Moderate”, and clinical records of absence of cognitive impairment), and Control (Braak 0-I, CERAD “None” or “Sparse”, and clinical records of absence of cognitive impairment). Subjects presenting comorbidities (frontotemporal lobar degeneration (FTLD), vascular dementia, Lewy body dementia, limbic-predominant age-related TDP-43 encephalopathy (LATE), and diabetes) were excluded. Blocks of formalin-fixed paraffin-embedded frontal lobe and mesial temporal lobe were sectioned (5 μm thickness) onto standard glass slides. Slides containing human brain tissue were processed for multiplex immunofluorescence and enzymatic immunohistochemistry.

### Multiplex immunofluorescence

Multiplex immunofluorescence (mIF) was achieved using the Opal 6-Plex Detection Kit (Akoya Biosciences, formerly Opal Polaris 7 Color IHC Automated Detection Kit). Staining was performed according to the manufacturer’s instructions. Frontal lobe sections, containing the DLPFC, were stained for β-tubulin (ab52623, Abcam, 1:200, 780 channel), GABA (PA5-32241, Thermo Fisher Scientific, 1:150, 690 channel), somatostatin (or SST, PA5-82678, Thermo Fisher Scientific, 1:200, channel 530), RBFOX1 (ab254413, Abcam, 1:200, channel 480), KIF26B (ab121952, Abcam, 1:100, channel 570), and Aβ (6E10, SIG-39300, BioLegend, 1:1000, channel 620). Mesial temporal lobe sections, containing the EC, were stained for β-tubulin (ab52623, Abcam, 1:200, 780 channel), NeuN (ab177487, Abcam, 1:150, channel 690), Reelin (or RELN, ab312310, Abcam, 1:150, channel 520), ATP8B1 (PA5-53839, Thermo Fisher Scientific, 1:150, channel 480), MEF2C (ab211493, Abcam, 1:150, channel 570), and Aβ (6E10, SIG-39300, BioLegend, 1:1000, channel 620). Tissue sections were imaged at 40x magnification on a PhenoImager HT (Akoya Biosciences) using whole slide scan settings. Brightness and contrast were adjusted using QuPath (version 0.4.3). Image processing was performed using *QuPath* [124], with cell detection based on DAPI being achieved using StarDist. Cell quantification was accomplished using a machine learning (ML) classifier within *QuPath*.

#### Analysis of single-cell protein quantifications from Qupath

Raw quantifications of marker intensities of 16 DLPFC samples (Control=6, AD=6, Resilient=4) and 15 EC samples (Control=6, AD=6, Resilient=5) were analyzed independently by region. Qupath quantifications were filtered first based on the upper and lower 1% percentiles of cell area, cell length and nucleus diameter; this approach is similar to that followed in [28]. Subsequently, cells with autofluorescence values below the 1% percentile or higher than the 5% percentile were discarded from the analysis. In addition, cells from the EC were also filtered based on the mean intensity of NeuN, following an approach similar to that taken in single cell transcriptomic data to discard cells with high mitochondrial content. After filtering, each dataset was analyzed following the protocol for image-based spatial data analysis in Seurat 5.1.0 [125]. The subcellular organization of protein markers was preserved in downstream analysis by taking the mean intensity of each marker in each of three cellular compartments (nucleus, cytoplasm and membrane), and treating each compartment as an independent variable. For example, MEF2C quantifications resulted in three variables: MEF2C-Nucleus, MEF2C-Membrane and MEF2C-Cytoplasm. The spatially resolved data was imported into Seurat using the function *LoadAkoya.* The dataset was normalized and scaled using the centered log-ratio method, as recommended by the protocol. Principal component analysis was used to reduce the number of variables in each dataset by 50%. A UMAP embedding was obtained from the resulting principal components. The same number of principal components were used to find neighbors and perform clustering with a resolution of 0.4 in both datasets. Clusters were manually annotated by inspecting the expression of all the protein targets measured in each brain region. Cell types were named according to their RNA-derived counterparts whenever possible.

#### Comparison of cell proportions using protein-derived cell types

Once individual cell types were annotated, the comparison of cellular proportions was done following the same procedure described for the transcriptomic-based comparison, with a few modifications. The cellular proportions in the EC datasets are based on the pool of NeuN+ cells, which were removed from the beginning of the analysis with Seurat. In the DLPFC dataset, cells were not pre-filtered based on a neuronal marker; this was done after clustering. The population labeled as GABA-cells was discarded before the analysis of proportions. We opted for this approach due to the empirical distribution of GABA, which as opposed to the distribution of NeuN in the EC, had no obvious cutoff to split the neuronal population.

#### Analysis of marker co-expression

Co-expression of selected markers and their association with phenotypic data (e.g., disease group) were investigated using the processed mIF DLPFC and EC datasets separately. First, we calculated Pearson correlations between all pairwise combinations of markers in every cell population using the normalized intensities from Seurat. The resulting P values were adjusted for multiple testing using the FDR method. For additional inspection of co-expression, an alternative series of linear models were used to account for additional covariates (general design formula: *response marker ∼ predictor marker + cell type + age + sex + disease group*) These formulations allow for reporting adjusted R-squared values, corresponding to the co-expression between the markers, as well as their associated P values.

### Polygenic risk scores

To examine the genetic risk underlying each of the diagnostic groups, polygenic risk scores (PRS) were calculated for each subject from ROSMAP with genetic data available [20] (n = 986) individually with PRSice-2 [126]using published effect sizes for 85 SNPs [9]. Each subject was classified as AD (n = 574), RES (n = 304), or CTRL (n = 108), as described for the bulk RNAseq analysis. Odds ratios were taken from published GWAS data [9] and applied to the samples used here to get individual PRS, and then compared across groups. All PRS were z-score normalized. A student’s T-test was used to compare sample groups with correction for multiple testing.

## Declarations

### Availability of data and materials

The datasets analyzed for the current study are available in Synapse. Bulk RNAseq data and snRNAseq data are accessible under the accession codes from their original publications syn8456629 and syn52293442, respectively, under controlled use conditions due to human privacy regulations. The multiplex immunofluorescence data along with the results for all analyses have been deposited on Synapse under the Synapse ID <u>syn63686123</u>. All code used for this publication is available at https://github.com/hidelab/ADresilience_CastanhoNaderi.git.

### Author’s contributions

I.C., P.N., and W.H. designed the study. W.H. and I.C. secured funding. S.M. generated the polygenic risk scores. P.N. downloaded, processed, and analyzed the bulk RNAseq data. H.M. and L.H.T. generated the snRNAseq data, and C.B. and H.M. preprocessed it. I.C. processed and analyzed the snRNAseq. D.P. generated and processed the whole genome sequencing data. B.W. collected the clinical information for the BIDMC cohort and assessed the neuropathological phenotypes for each subject in the cohort. A.P. generated the multiplex immunofluorescence data at the Spatial Technologies (STU) core at BIDMC, I.C. processed it with support from N.K., and P.N. and L.S. performed the analysis. I.C., P.N., and W.H. interpreted the results, with input from R.T and D.Y.L, and feedback from all co-authors. I.C., P.N., and W.H. wrote the manuscript, which was approved by all authors. D.A.B. is PI of ROS and MAP and oversaw all of the clinical and neuropathologic data collection and biospecimen distributions for omics, and critically reviewed the manuscript.

## Acknowledgements

We thank all investigators from the CIRCUITS consortium (Collaboration to Infer Regulatory Circuits and Uncover Innovative Therapeutic Strategies consortium) and AGP (Alzheimer’s Genome Project) kindly funded by the Cure Alzheimer’s Fund (CureAlz). We acknowledge Spatial Technologies Unit of Precision RNA Medicine Core (RRID:SCR_024905) and thank Antonella de Amaral and Shuoshuo Wang for their technical assistance. We also thank Yered Pita-Juarez, Dimitra Karagkouni, and Ioannis Vlachos for important input into discussion of bioinformatics methods. We thank Drs. Quadri Adewale and Lester Kobzik for their insights and proofreading of the manuscript. Figures 1A, 3A, 4A, 5, and S1A were created with BioRender.com. Figure 5 was adapted from the templates “Excitatory and inhibitory neurons distribution in the layers of the human cortex”, “Cleavage of Amyloid Precursor Protein (APP)”, and “Alzheimer’s Brain (Disintegrating Microtubule)” by BioRender.com (2024). Retrieved from https://app.biorender.com/biorender-templates.

## Supplementary figures

**Figure S1.**
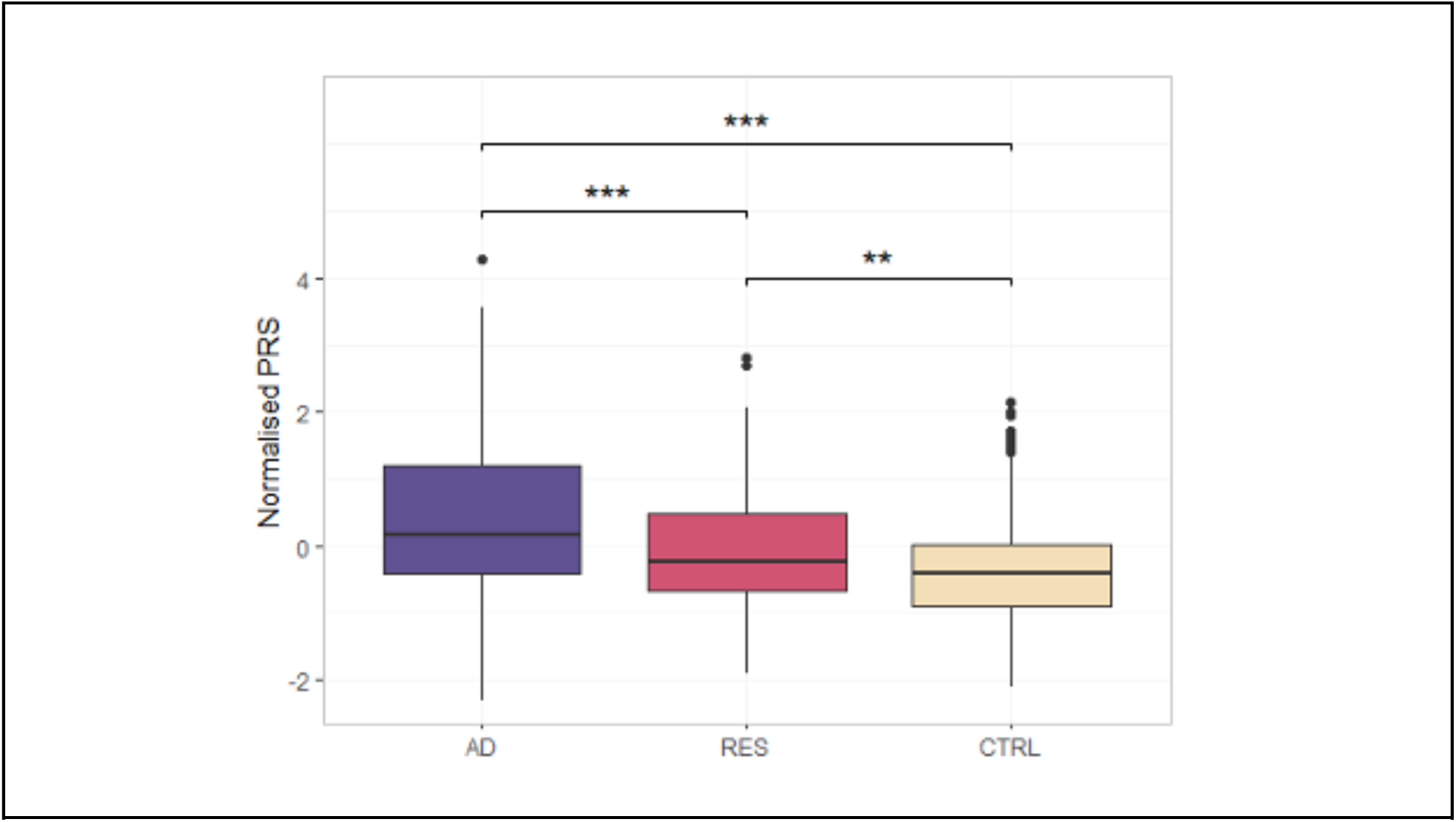
Sample classification details and cognitive resilience relative to AD polygenic risk. **(A)** AD polygenic risk scores (AD-PRS) were calculated for each subject from ROSMAP with genetic data available. The central horizontal line of the box plots depicts the median, and the lower and upper hinges correspond to the first and third quartiles (the 25th and 75^th^ percentiles). The circles represent outliers. Bonferroni adjusted P values. ** Adj-P < 0.01, *** Adj-P < 0.001. CTRL: Control, AD: Alzheimer’s disease, RES: Resilient.

**Figure S2.**
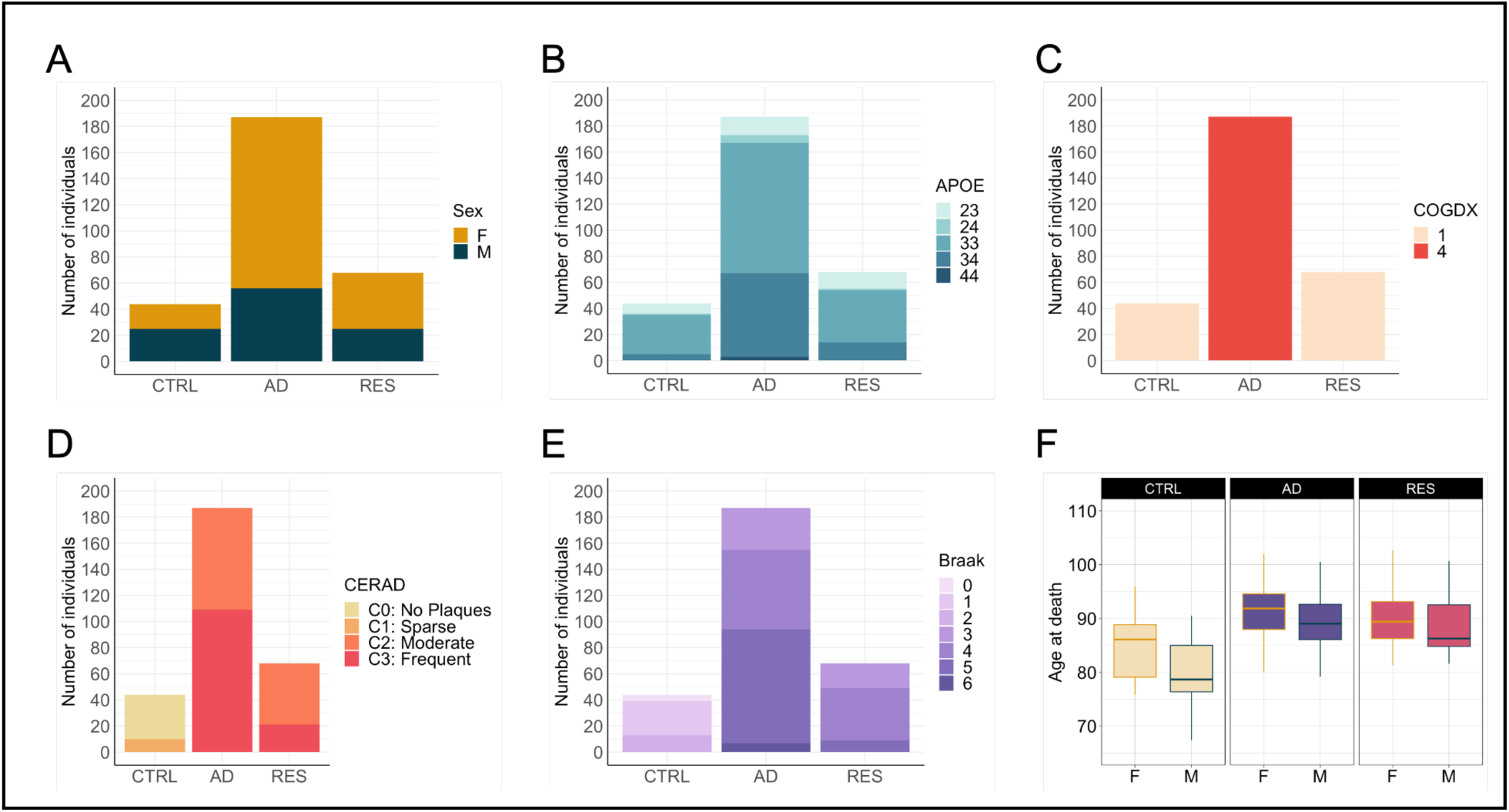
Characteristics of bulk RNAseq ROSMAP subjects used in this study (DLPFC). **(A)** Sex distribution. **(B)** *APOE* genotype distribution. **(C)** Clinical consensus diagnosis of cognitive status at the time of death (final consensus cognitive diagnosis, “cogdx” variable from RUSH Alzheimer’s Disease Center (RADC) Research Resource Sharing Hub). **(D)** Consortium to Establish a Registry for Alzheimer’s Disease (CERAD) score. **(E)** Braak stage distribution. **(F)** Age distribution, by sex. The central horizontal line of the box plots depicts the median, and the lower and upper hinges correspond to the first and third quartiles (the 25th and 75th percentiles). CTRL: Control, AD: Alzheimer’s disease, RES: Resilient. F: females, M: males.

**Figure S3.**
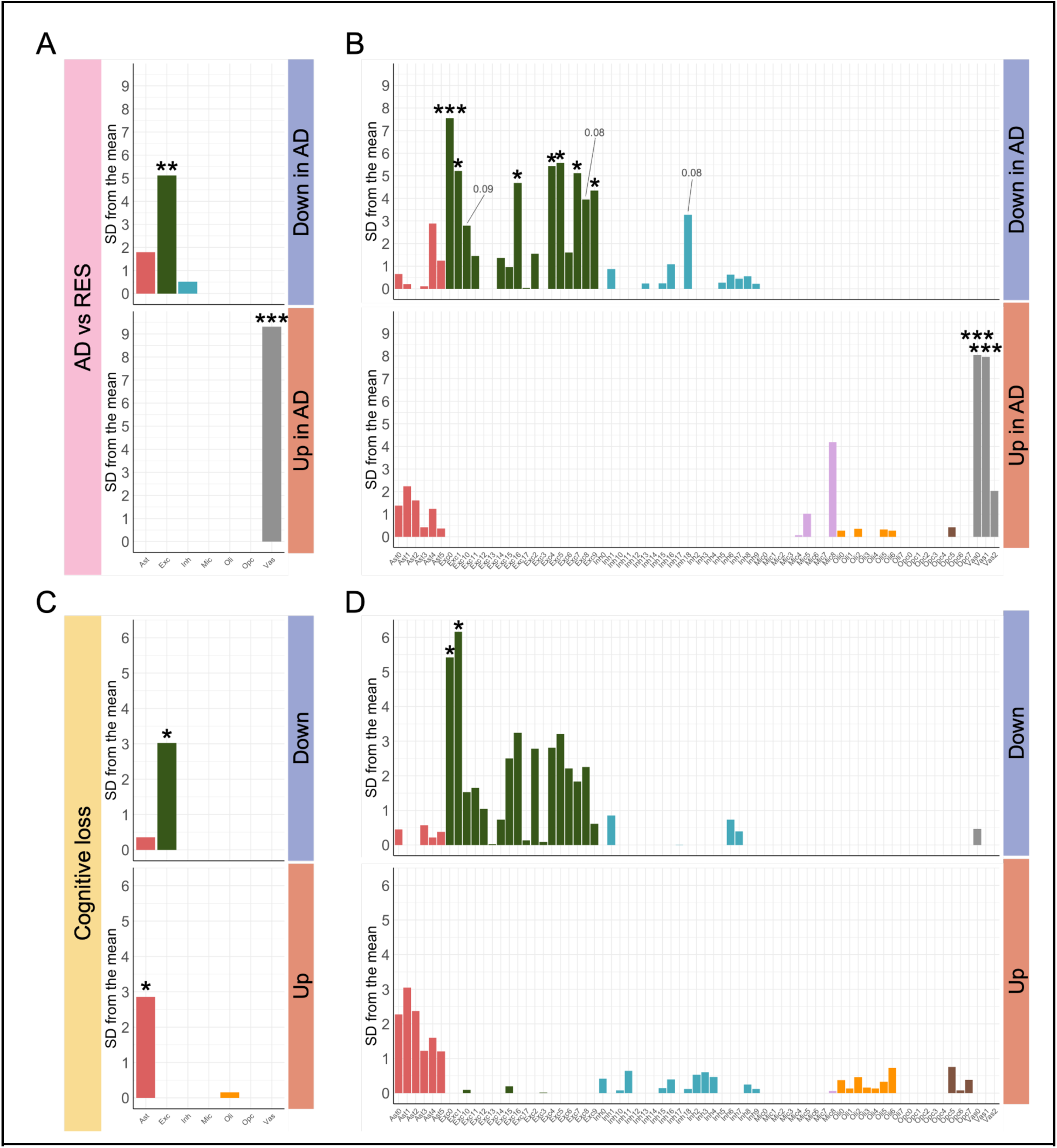
Expression weighted cellular enrichment results for differentially expressed genes in bulk RNAseq data from DLPFC. (A-B) Cellular enrichment of genes uniquely differentially expressed in ADvsRES (61 DEGs) in major cell types (A) and cell subtypes (B) identified in the DLPFC. (B-C) Cellular enrichment for genes associated with cognitive decline in major cell types (A) and cell subtypes (B). Down (top): genes down-regulated with cognitive decline; Up (bottom): genes up-regulated with cognitive decline. Y-axis shows the number of standard deviations from the bootstrap mean, associated with the enrichment of the stated cell type in the gene list investigated. Stars represent adjusted P < 0.05 (Bonferroni correction).

**Figure S4.**
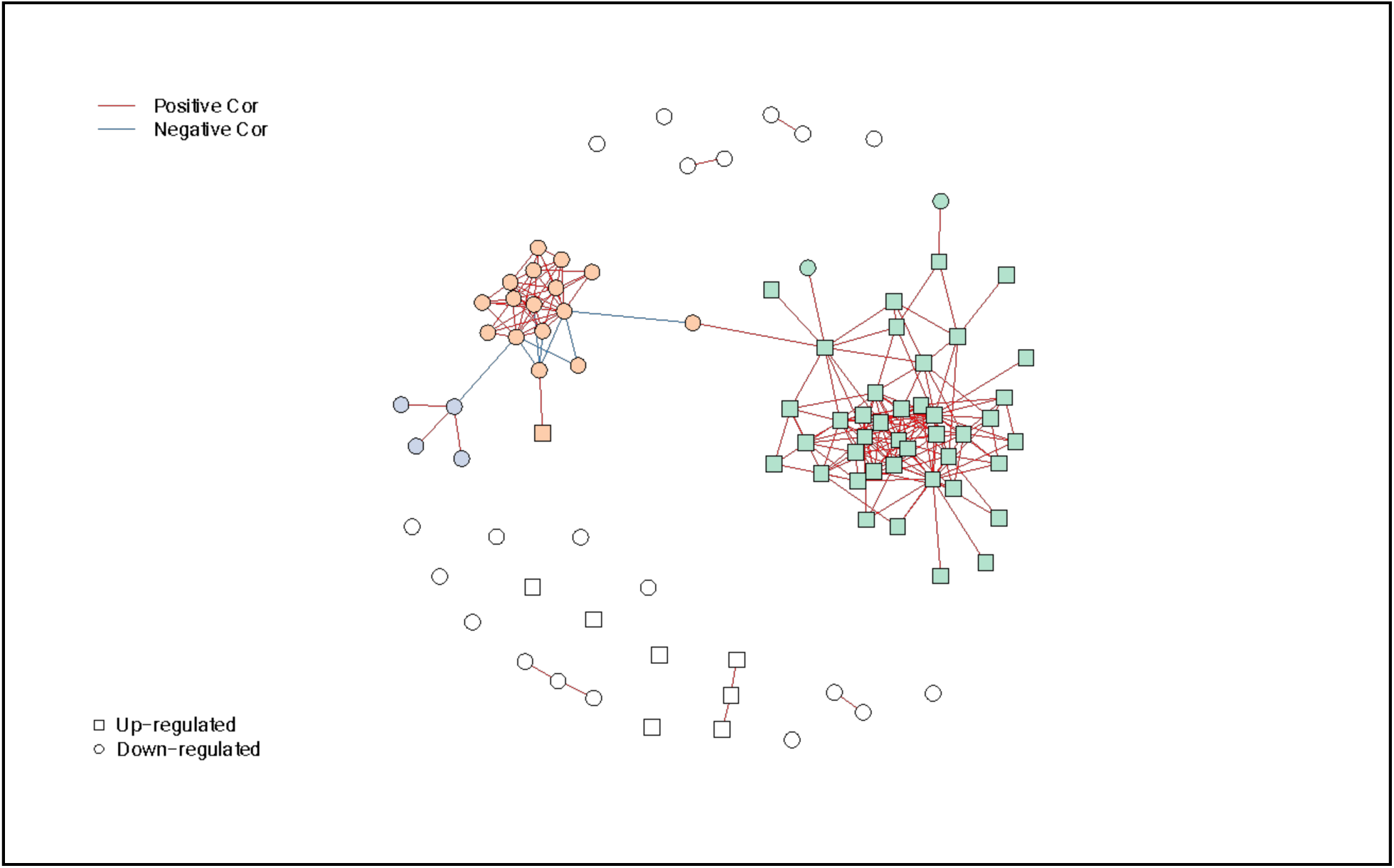
Resulting clusters from pathway activity analysis for ADvsRES. Nodes represent pathways with significant dysregulation of activity (q-value < 0.1) in ADvsRES. Out of a total of 99 dysregulated pathways between ADvsRES, 62 pathways organized in two unsupervised clusters of expression. Node shapes denote up-regulation and down-regulation in ADvsRES. Edges represent canonical co-expression of pathways based on the Pathway Co-expression Network background. Pathway activity profiles were determined using the PanomiR software package. Pathway dysregulation p-values were determined using the Limma package’s linear regression models contrasting between ADvsRES and accounting for confounding covariates such as age, batch, and RNA integrity number.

**Figure S5.**
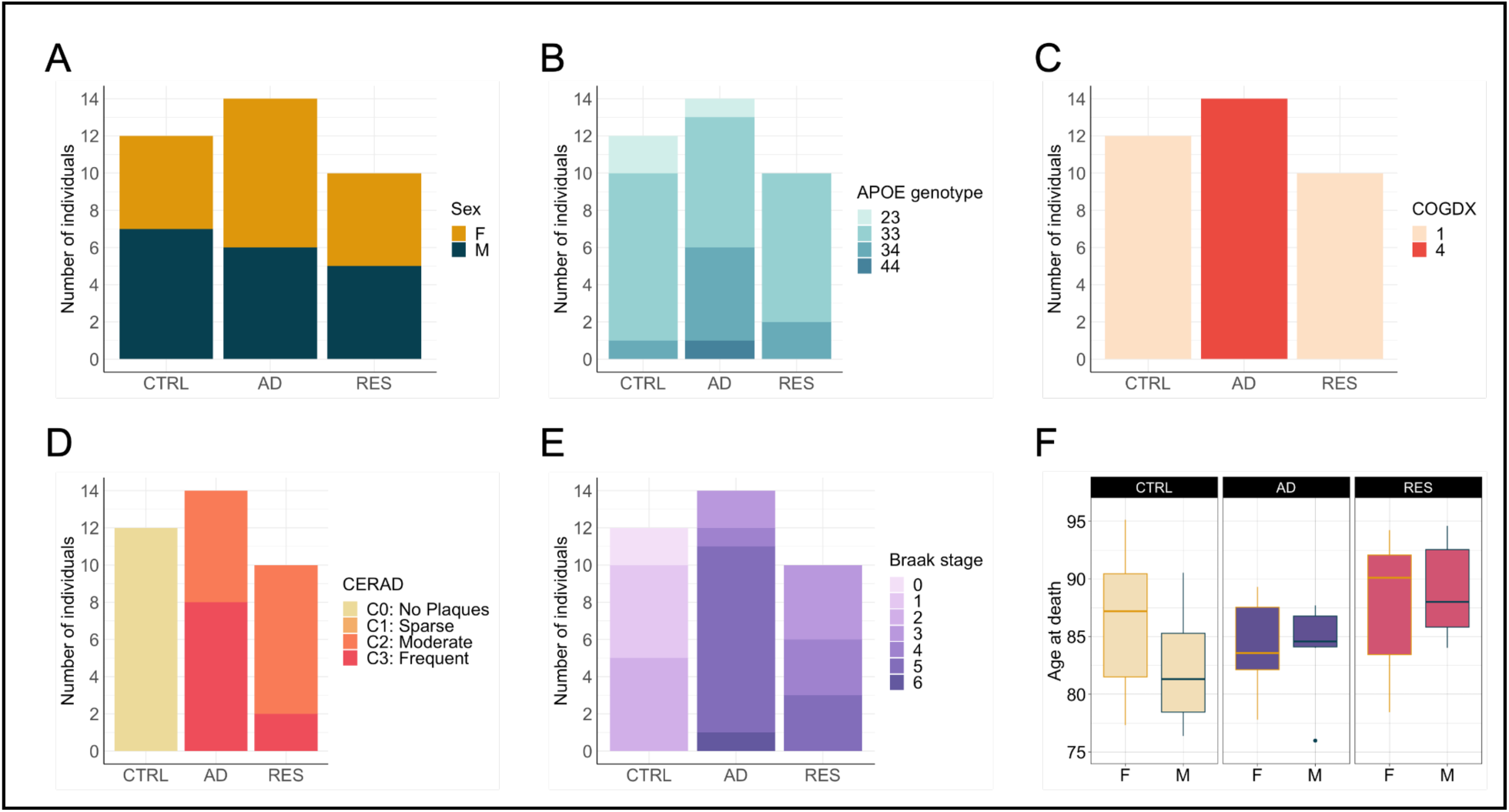
Characteristics of ROSMAP subjects with snRNAseq from the dorsolateral prefrontal cortex (DLPFC) used in this study. **(A)** Sex distribution. **(B)** APOE genotype distribution. **(C)** Clinical consensus diagnosis of cognitive status at time of death (final consensus cognitive diagnosis, “cogdx” variable from RUSH Alzheimer’s Disease Center (RADC) Research Resource Sharing Hub). **(D)** Consortium to Establish a Registry for Alzheimer’s Disease (CERAD) score. **(E)** Braak stage distribution. **(F)** Age distribution, by sex. The central horizontal line of the box plots depicts the median, and the lower and upper hinges correspond to the first and third quartiles (the 25th and 75th percentiles). Circles represent outliers. CTRL: Control, AD: Alzheimer’s disease, RES: Resilient. F: females, M: males.

**Figure S6.**
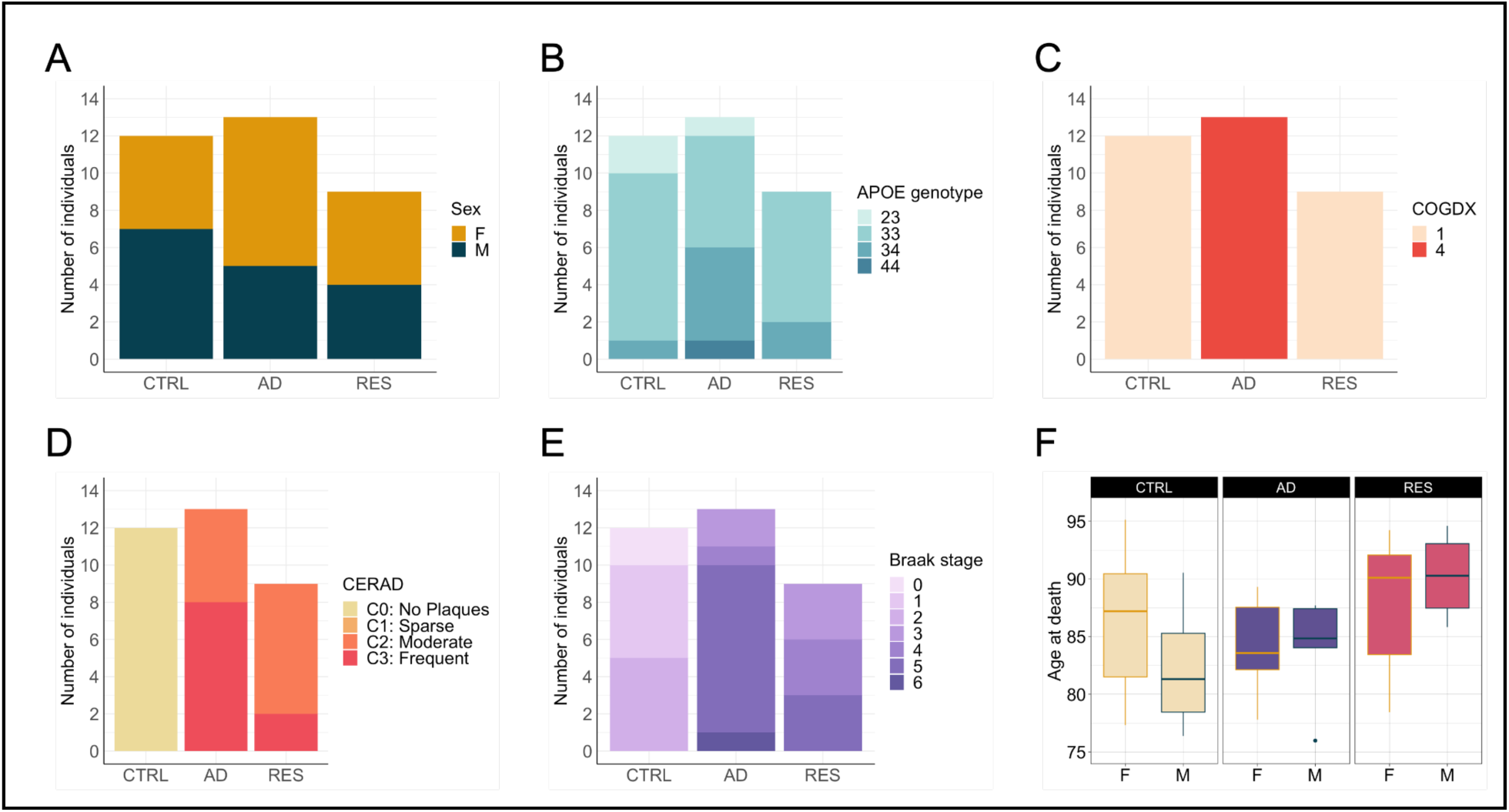
Characteristics of ROSMAP subjects with snRNAseq from the entorhinal cortex (EC) used in this study. **(A)** Sex distribution. **(B)** APOE genotype distribution. **(C)** Clinical consensus diagnosis of cognitive status at time of death (final consensus cognitive diagnosis, “cogdx” variable from RUSH Alzheimer’s Disease Center (RADC) Research Resource Sharing Hub). **(D)** Consortium to Establish a Registry for Alzheimer’s Disease (CERAD) score. **(E)** Braak stage distribution. **(F)** Age distribution, by sex. The central horizontal line of the box plots depicts the median, and the lower and upper hinges correspond to the first and third quartiles (the 25th and 75th percentiles). Circles represent outliers. CTRL: Control, AD: Alzheimer’s disease, RES: Resilient. F: females, M: males.

**Figure S7.**
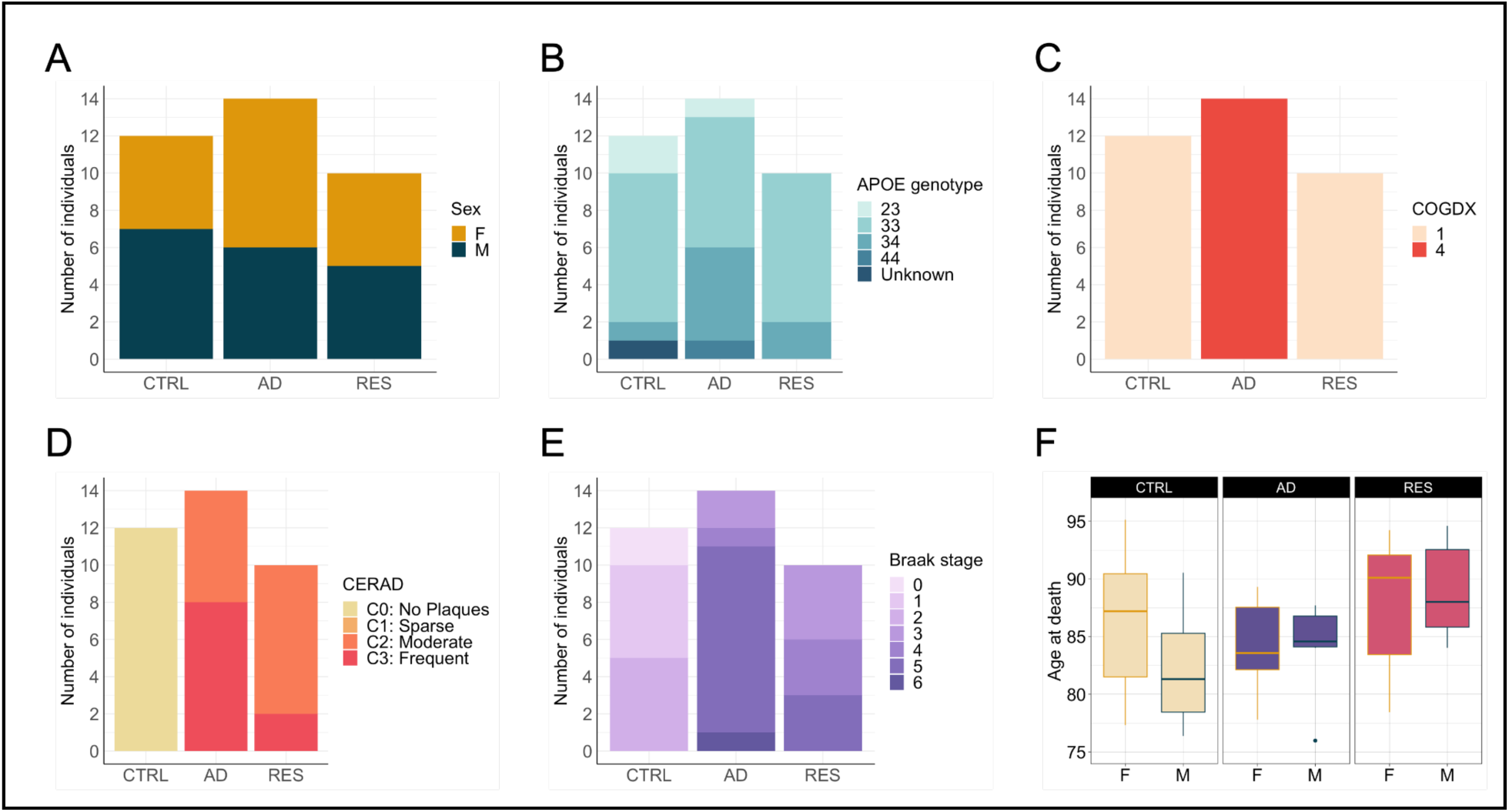
Characteristics of ROSMAP subjects with snRNAseq from the hippocampus (HC) used in this study. **(A)** Sex distribution. **(B)** APOE genotype distribution. **(C)** Clinical consensus diagnosis of cognitive status at time of death (final consensus cognitive diagnosis, “cogdx” variable from RUSH Alzheimer’s Disease Center (RADC) Research Resource Sharing Hub). **(D)** Consortium to Establish a Registry for Alzheimer’s Disease (CERAD) score. **(E)** Braak stage distribution. **(F)** Age distribution, by sex. The central horizontal line of the box plots depicts the median, and the lower and upper hinges correspond to the first and third quartiles (the 25th and 75th percentiles). Circles represent outliers. CTRL: Control, AD: Alzheimer’s disease, RES: Resilient. F: females, M: males.

**Figure S8.**
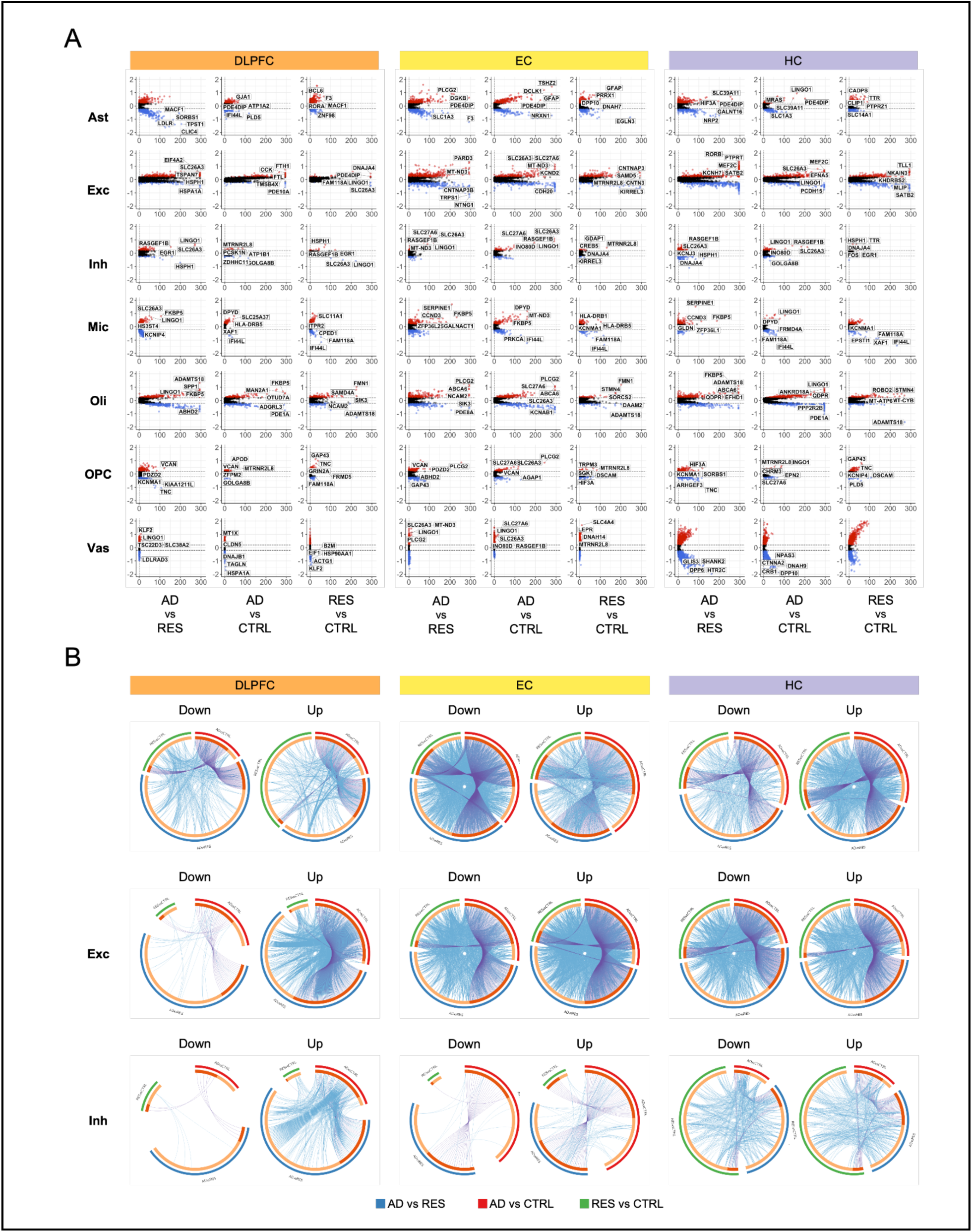
Cell-specific transcriptomic changes in AD resilience. **(A)** Volcano plots showing significantly (Bonferroni-corrected P < 0.1) differentially expressed genes (DEGs) in each brain region tested (DLPFC, EC, and HC) in ADvsRES, ADvsCTRL, and RESvsCTRL. DEGs with log2FC < −0.2 are highlighted in blue, and DEGs with log2FC > 0.2 are highlighted in red. The horizontal lines represent Bonferroni-adjusted P = 0.1. **(B)** Circular plots showing overlapping genes and ontologies from each comparison for excitatory and inhibitory neurons from each brain region investigated. Each outer arc represents the identity of each gene list (blue: ADvsRES, red: ADvsCTRL, green: RESvsCTRL). Each inner arc shows the genes that are shared by multiple lists in dark orange and genes that are unique to that gene list in light orange. Purple lines link the same gene, shared by multiple lists. Blue lines represent genes that fall under the same ontology term. Gene ontology enrichment analysis was performed using Metascape. CTRL: Control, AD: Alzheimer’s disease, RES: Resilient. Ast: Astrocytes, Exc: Excitatory neurons, Inh: Inhibitory neurons, Mic & Imm: Microglia and immune cells, Oli: Oligodendrocytes, OPC: Oligodendrocyte progenitor cells, Vas & Epi: Vascular and epithelial cells. DLPFC: Dorsolateral prefrontal cortex, EC: Entorhinal cortex, HC: Hippocampus.

**Figure S9.**
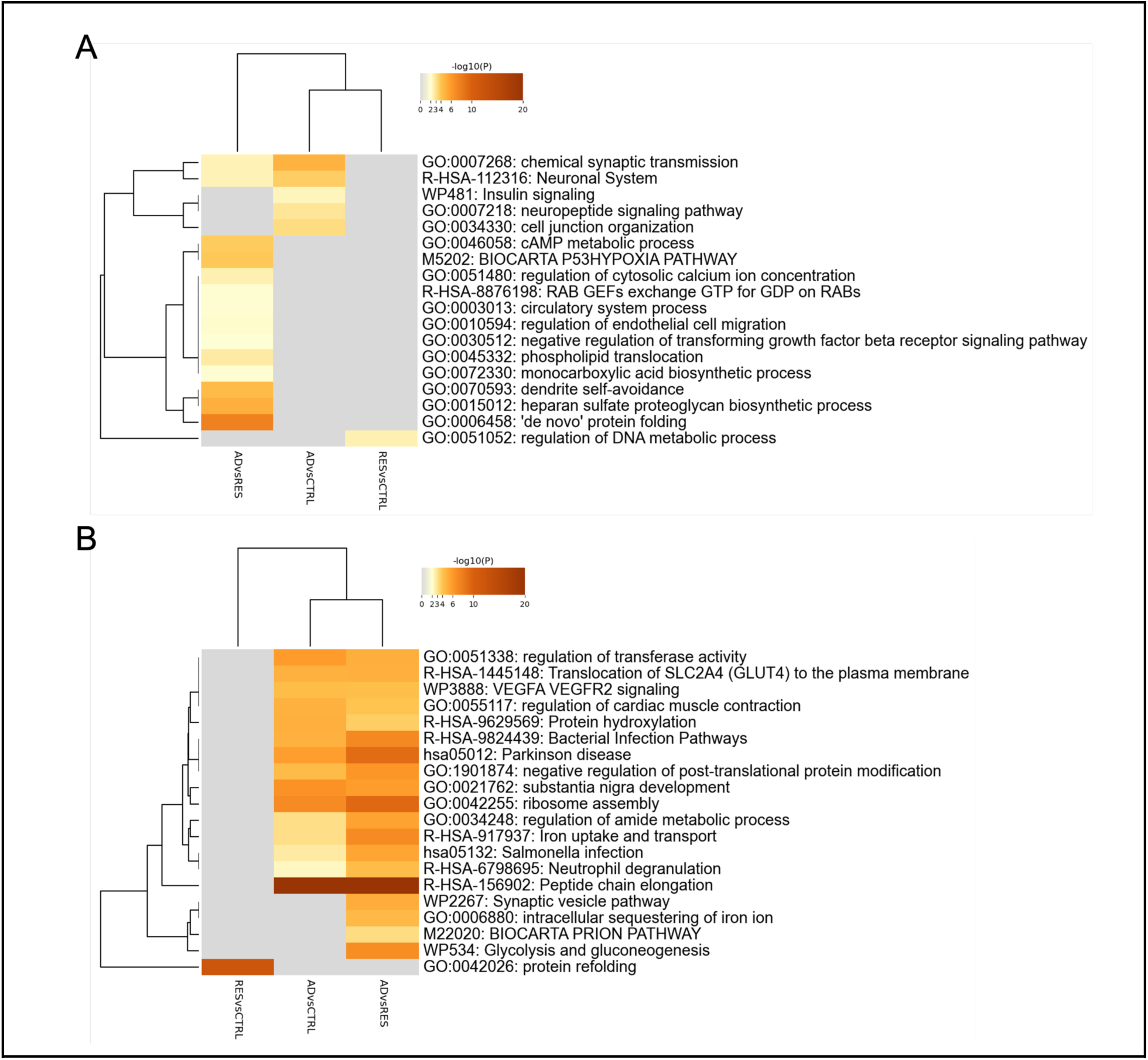
Significant terms from gene ontology enrichment analysis for DEGs identified in excitatory neurons from the DLPFC. **(A-B)** Heatmaps showing enriched gene ontology clusters across gene lists from each of the three comparisons (ADvsRES, ADvsCTRL, RESvsCTRL). The cells in each heatmap are colored by their respective p-values, with gray cells indicating a lack of enrichment for that term in the corresponding gene list. The terms with the best p-values within each cluster are displayed in the dendrogram. Cumulative hypergeometric p-values and enrichment factors were calculated and used for filtering. Significant terms were hierarchically clustered into a tree based on Kappa-statistical similarities among their gene memberships, with 0.3 kappa score applied as the threshold to cast the tree into clusters. **(A)** Genes down-regulated in the first group compared to the second group for each comparison. **(B)** Genes up-regulated in the first group compared to the second group for each comparison.

**Figure S10.**
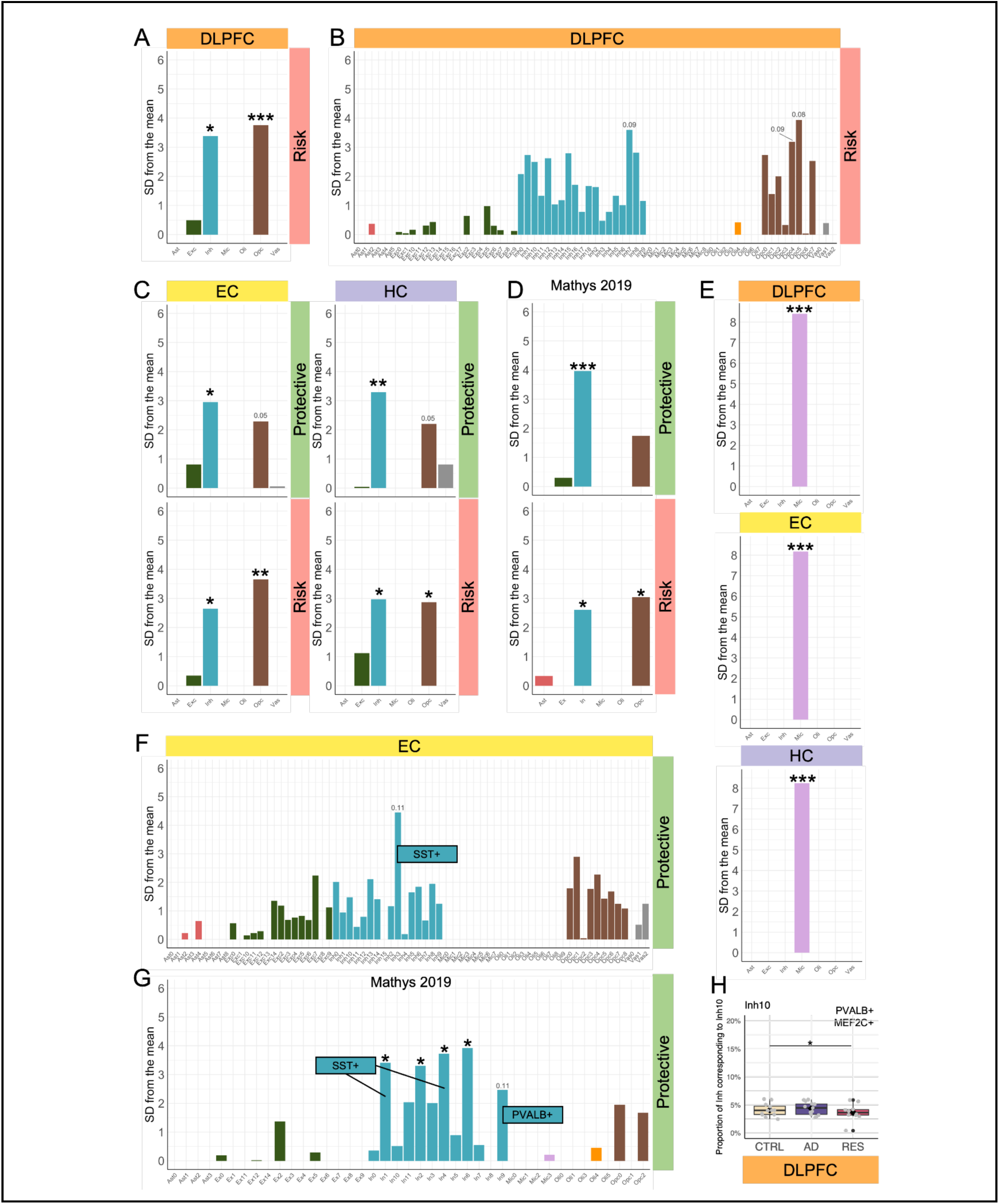
Expression-weighted cellular enrichment results for genes identified from rare variants. **(A)** Cellular enrichment of genes identified from risk genetic rare variants in the DLPFC for major cell types. **(B)** Cellular enrichment of genes identified from risk genetic rare variants in the DLPFC for cell subtypes. **(C)** Cellular enrichment of genes identified from protective and risk genetic rare variants in the EC and HC for major cell types. **(D)** Cellular enrichment of genes identified from protective and risk genetic rare variants in the Mathys et al. 2019 dataset (DLPC) for major cell types. **(E)** Genes identified from common variants from Bellenguez et al. 2022 in major cell types. **(F)** Cellular enrichment of genes identified from protective rare variants in the EC for cell subtypes. **(G)** Cellular enrichment of genes identified from protective rare variants in the Mathys et al. 2019 dataset (DLPC) for cell subtypes. **(E)** Cell proportion changes for PVALB+ DLPFC:Inh10. * Adj-P < 0.05, ** Adj-P < 0.01, *** Adj-P < 0.001 CTRL: Control, AD: Alzheimer’s disease, RES: Resilient. DLPFC: Dorsolateral prefrontal cortex, EC: Entorhinal cortex, HC: Hippocampus.

**Figure S11.**
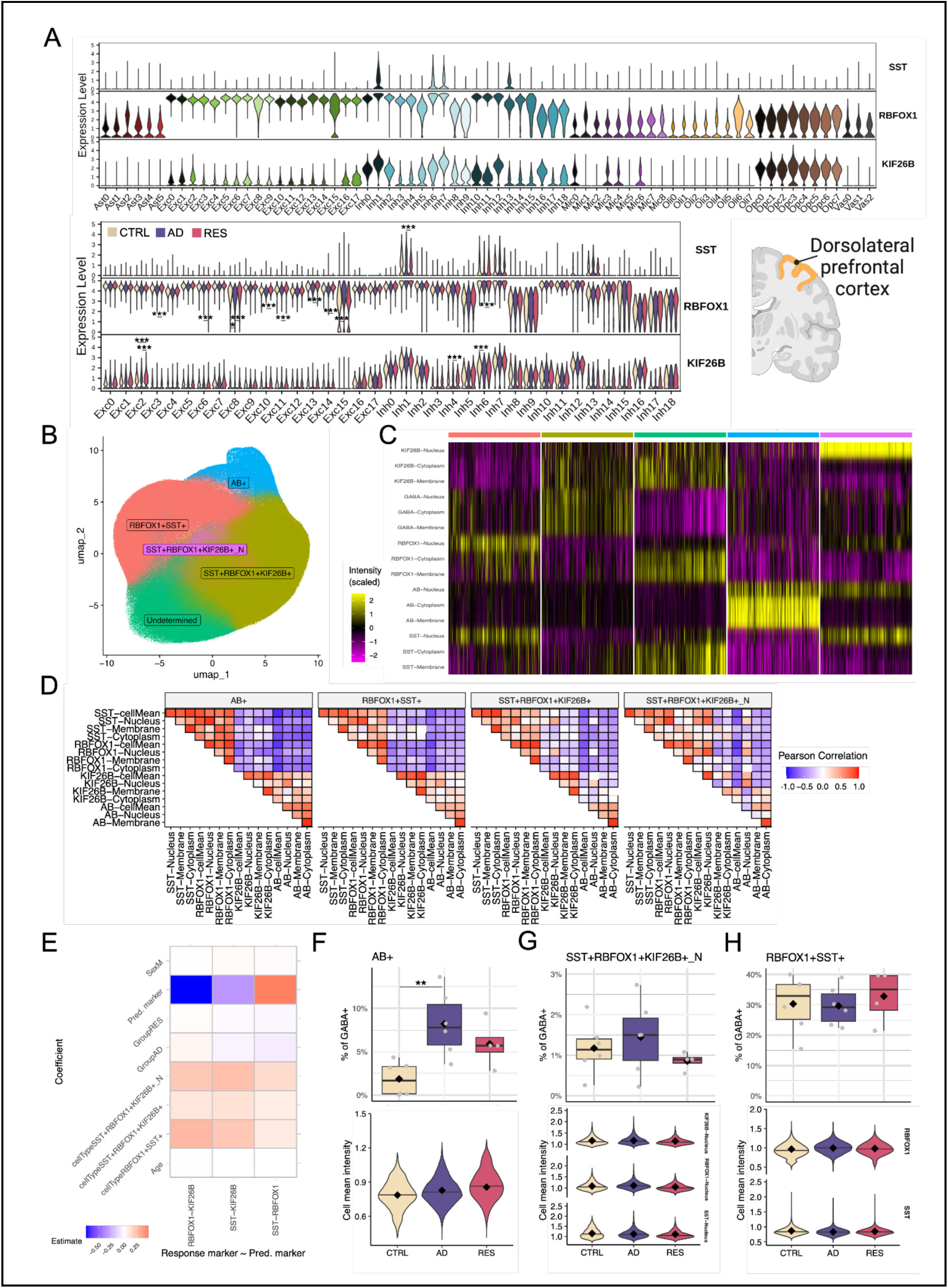
Gene expression distributions and protein expression assessed by mIF for SST, RBFOX1, and KIF26B. **(A)** Violin plots showing gene expression levels of SST, RBFOX1, and KIF26B for each subtype (top) and neuronal subtypes by diagnostic group (bottom) in the DLPFC (related to Figure 3L). **(B)** UMAP showing the populations identified in the DLPFC in the BIDMC cohort from multiplex immunofluorescence (mIF) staining for GABA, SST, RBFOX1, KIF26B, and Aβ. N_subjects_ = 16 (6 CTRL, 6 AD, 4 RES), N_cells_ = 1310803 (CTRL = 465007, AD = 559794, RES = 286002 cells). **(C)** Heatmap of normalized intensity of protein markers in a random subset of 500 cells from each cell type (indicated by the top color bar). Cell type labels shown in (B). **(D)** Correlation of inhibitory neuronal targets normalized intensity across identified cell subpopulations. Each square is colored by the Pearson correlation coefficient corresponding to the pair of markers indicated in the x and y axes. Solid black squares indicate significant correlations at an FDR of 0.05. **(E)** Coefficients of predictors used in linear models of marker intensities. **(F-H)** Box plots (top) showing the results for differential cell proportions of clusters shown in **(B)** and violin plots (bottom) showing the distribution of their corresponding cell-level protein intensities (normalized immunofluorescence levels) in each population. Diamonds show the grand mean of subject-level mean normalized intensities across cells. CTRL: Control, AD: Alzheimer’s disease, RES: Resilient. DLPFC: Dorsolateral prefrontal cortex. * Adj-P < 0.05, ** Adj-P < 0.01, *** Adj-P < 0.001.

**Figure S12.**
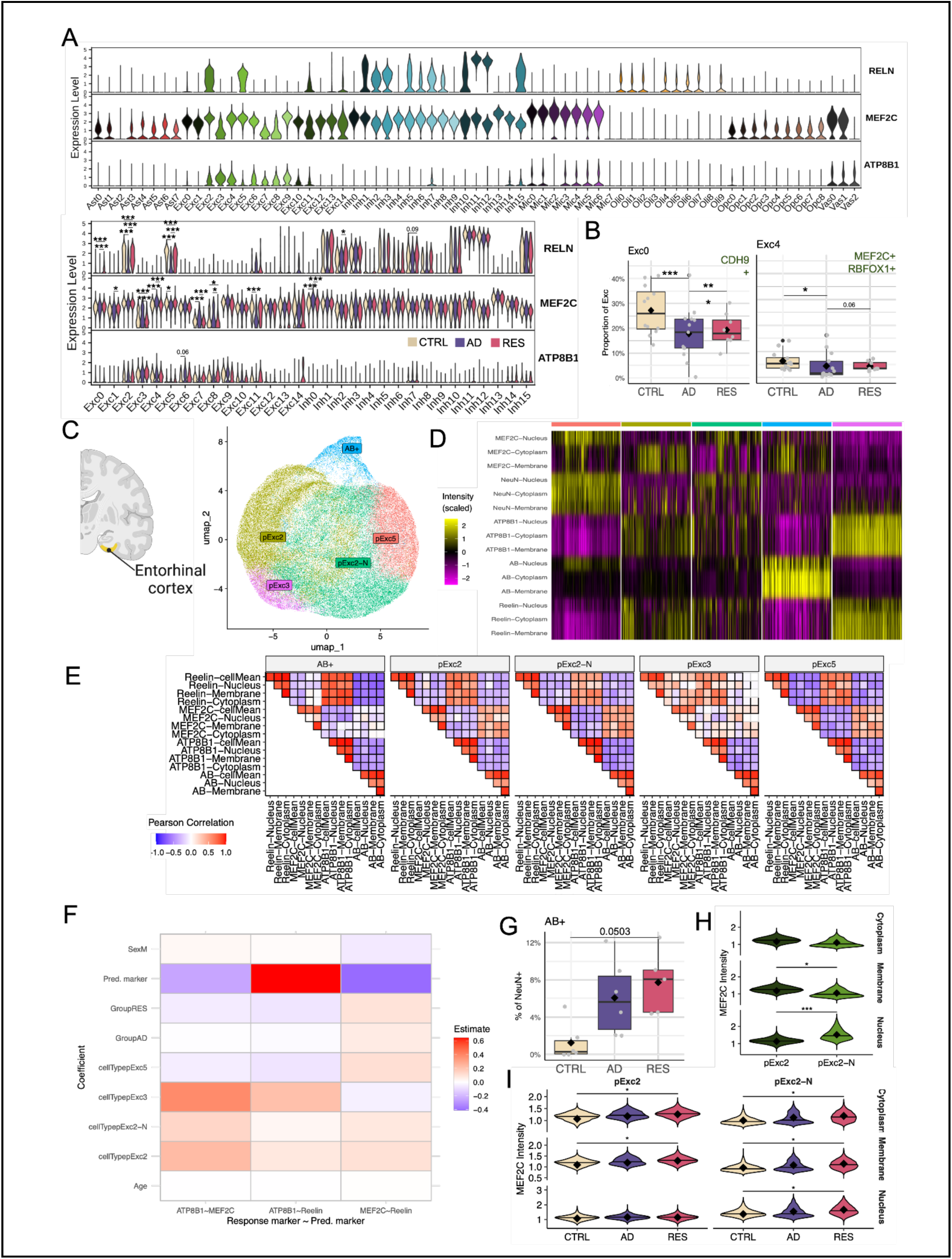
Significant cell proportion changes in vulnerable excitatory neurons and gene expression dynamics in excitatory neurons associated with resilience. **(A)** Violin plots showing gene expression levels (snRNAseq) for RELN, MEF2C, and ATP8B1 for all subtypes (top) and neuronal subtypes by group (bottom) in the EC (related to Figure 4). Cell proportion changes in additional vulnerable excitatory subtypes from the entorhinal cortex, as reported by Leng et al., 2021. **(B)** Cell proportion changes for CDH9+ EC:Exc0 cells (left) and cell proportion changes for RBFOX1+ MEF2C_high_ EC:Exc4 (right) **(C)** UMAP showing the populations identified in the EC in the BIDMC cohort from multiplex immunofluorescence (mIF) staining for NeuN, MEF2C, ATP8B1, RELN, and Aβ. N_subjects_ = 16 (6 CTRL, 6 AD, 5 RES), N_cells_ = 81549 cells. **(D)** Heatmap of normalized intensity of protein markers in a random subset of 500 cells from each cell type (indicated by the top color bar). Cell type labels shown in **(B)**. **(E)** Correlation of inhibitory neuronal targets normalized intensity across identified cell subpopulations. Each square is colored by the Pearson correlation coefficient corresponding to the pair of markers indicated in the x and y axes. Solid black squares indicate significant correlations at an FDR of 0.05. **(F)** Coefficients of predictors used in linear models of marker intensities. **(G)** Box plots showing cell proportion changes in the cluster AB+ shown in **(B)**, corresponding to neurons expressing high protein levels of Aβ. **(H)** Violin plots showing protein expression (normalized immunofluorescence intensity levels) of MEF2C in different compartments (cytoplasm, membrane, and nucleus) for the clusters pExc2 and pExc2-N (MEF2Chigh ATP8B1+ RELN+) shown in **(D)**. **(I)** Violin plots showing distributions of protein expression (normalized immunofluorescence intensity levels) of MEF2C in different compartments (cytoplasm, membrane, and nucleus) for the clusters pExc2 and pExc2-N (MEF2Chigh ATP8B1+ RELN+) by diagnostic group (CTRL, AD, RES). In **(H)** and **(I)** stars indicate significance level based on nominal P values of a Wilcoxon test performed on the subject-level means. Diamonds show the grand mean of subject-level mean normalized intensities across cells. CTRL: Control, AD: Alzheimer’s disease, RES: Resilient. EC: Entorhinal cortex. * Adj-P < 0.05, ** Adj-P < 0.01, *** Adj-P < 0.001

**Figure S13.**
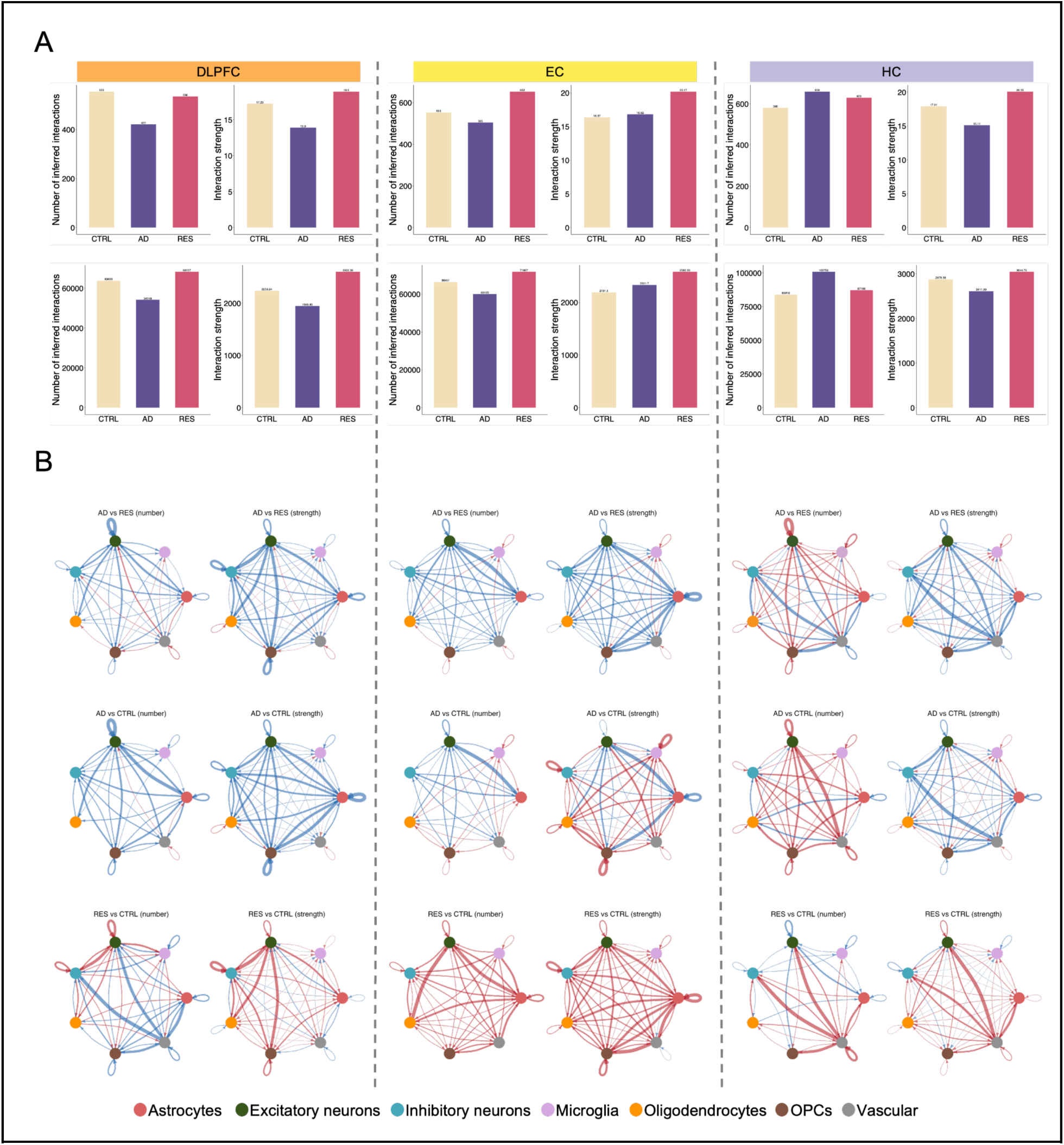
An increase in the number of events in cell-cell communication in resilience. **(A)** Bar plots showing the inferred number and strength of ligand-receptor interactions in each group for major cell populations (top) and cell subpopulations (bottom) in each brain region. **(B)** Differential number and strength of cell-cell interactions for each comparison. Arrows represent direction of interactions, from source to target major cell class. Blue represents a decrease in the number or strength of interactions between any two cells in the first compared to the second diagnostic group, and red represents an increase. CTRL: Control, AD: Alzheimer’s disease, RES: Resilient.

**Figure S14.**
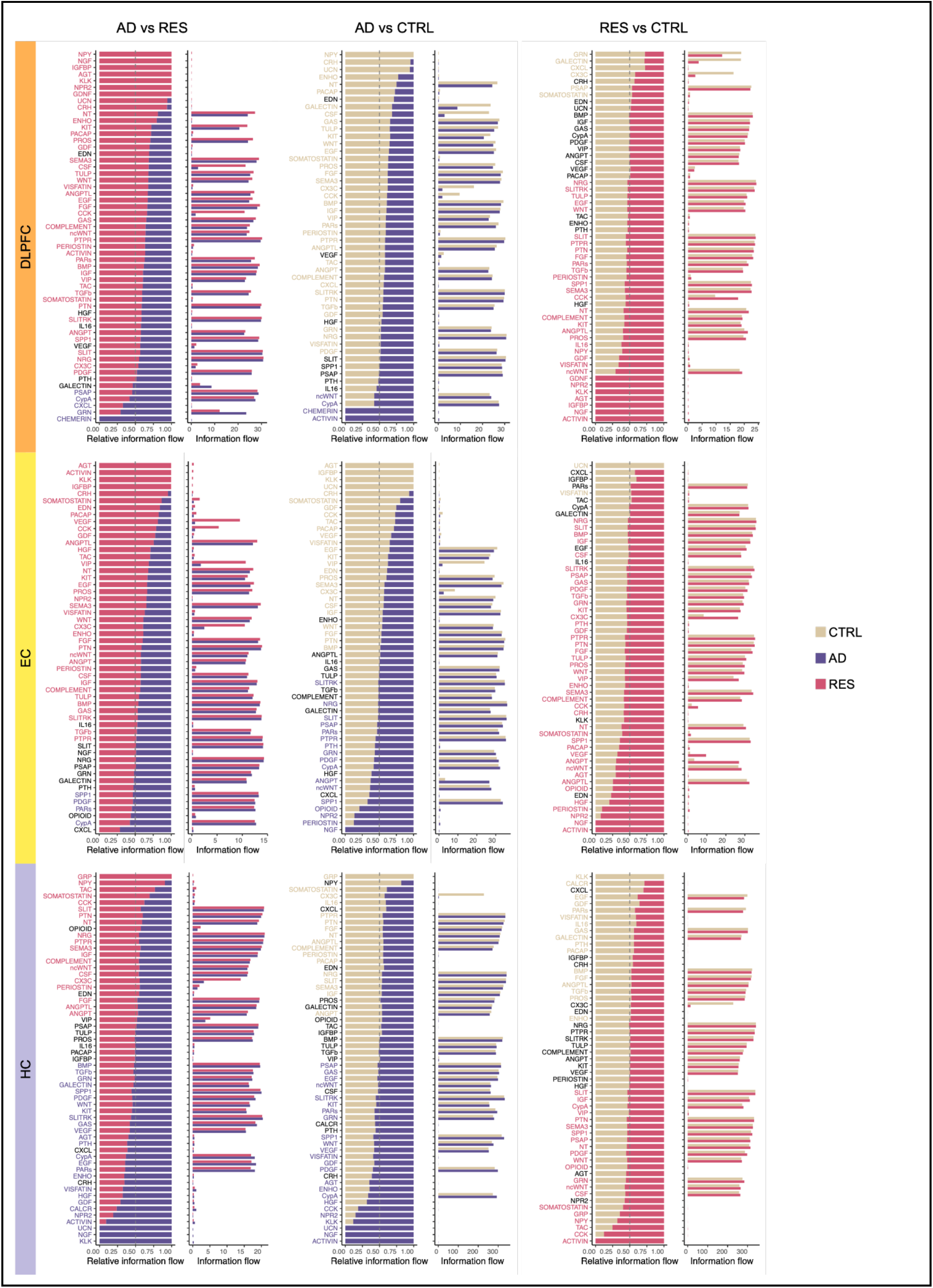
Significant pathways from intercellular communication analysis in cell subpopulations. List of significant pathways in each brain region (DLPFC, EC, and HC) for each comparison (ADvsRES, ADvsCTRL, and RESvsCTRL). Colored pathway names are significantly shifted towards the group with the corresponding color. Plots on the left show the relative information flow in one diagnostic group compared to the other, and the plots on the right show the overall information flow for each signaling pathway. CTRL: Control, AD: Alzheimer’s disease, RES: Resilient.

**Figure S15.**
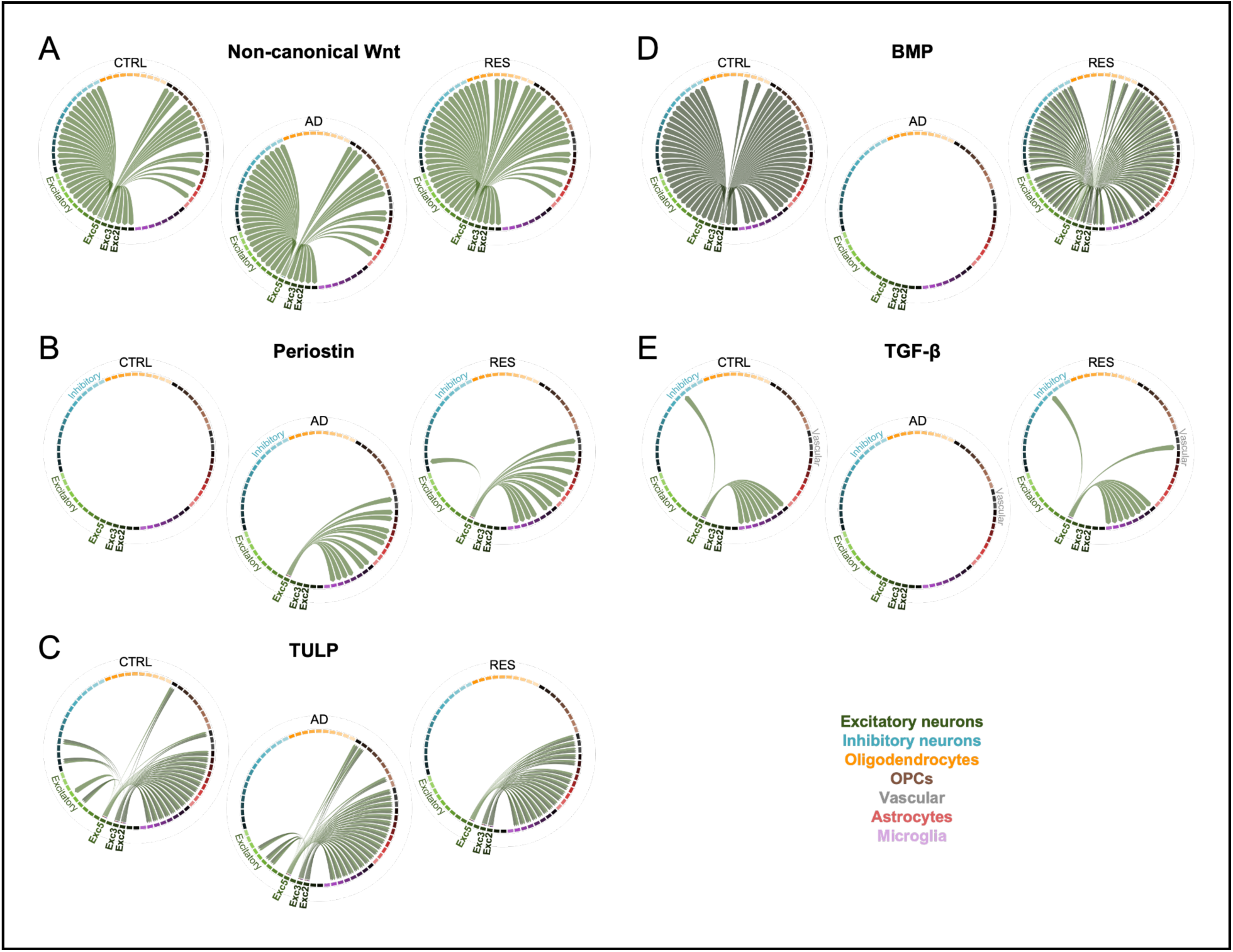
Signaling pathways emerging or disappearing in resilience in excitatory neuronal subpopulations from the entorhinal cortex showing a resilient phenotype. Chord diagrams showing significant networks (Figure S15) having EC:Exc2, EC:Exc:3, or EC:Exc5 as sources. **(A)** Non-canonical Wnt (source: EC:Exc5; targets: multiple subtypes of oligodendrocytes; ligand: WNT5A; receptor: MCAM). **(B)** Periostin (source: EC:Exc5; target: EC:Inh1; ligand: periostin; receptor: ITGAV/ITGB5). **(C)** TULP (sources: EC:Exc2, EC:Exc3, and EC:Exc5; targets: subtypes of excitatory neurons, inhibitory neurons, OPCs, and astrocytes; ligand: TUB; receptor: MERTK). **(D)** BMP (source: EC:Exc5; targets: subtypes from all major cell types; ligands: GDF7 and BMP8A; receptors: BMPR1A/ACVR2A, BMPR1A/ACVR2B, BMPR1A/BMPR2, BMPR1B/ACVR2A, BMPR1B/BMPR2, ACVR1/ACVR2A, ACVR1/BMPR2, BMPR1B/ACVR2B). **(E)** TGF-β (source: EC:Exc5; target: EC:Fib; ligand: TGF-β2; receptors: TGFBR1/R2 and ACVR1/TGFBR). CTRL: Control, AD: Alzheimer’s disease, RES: Resilient.

**Figure S16.**
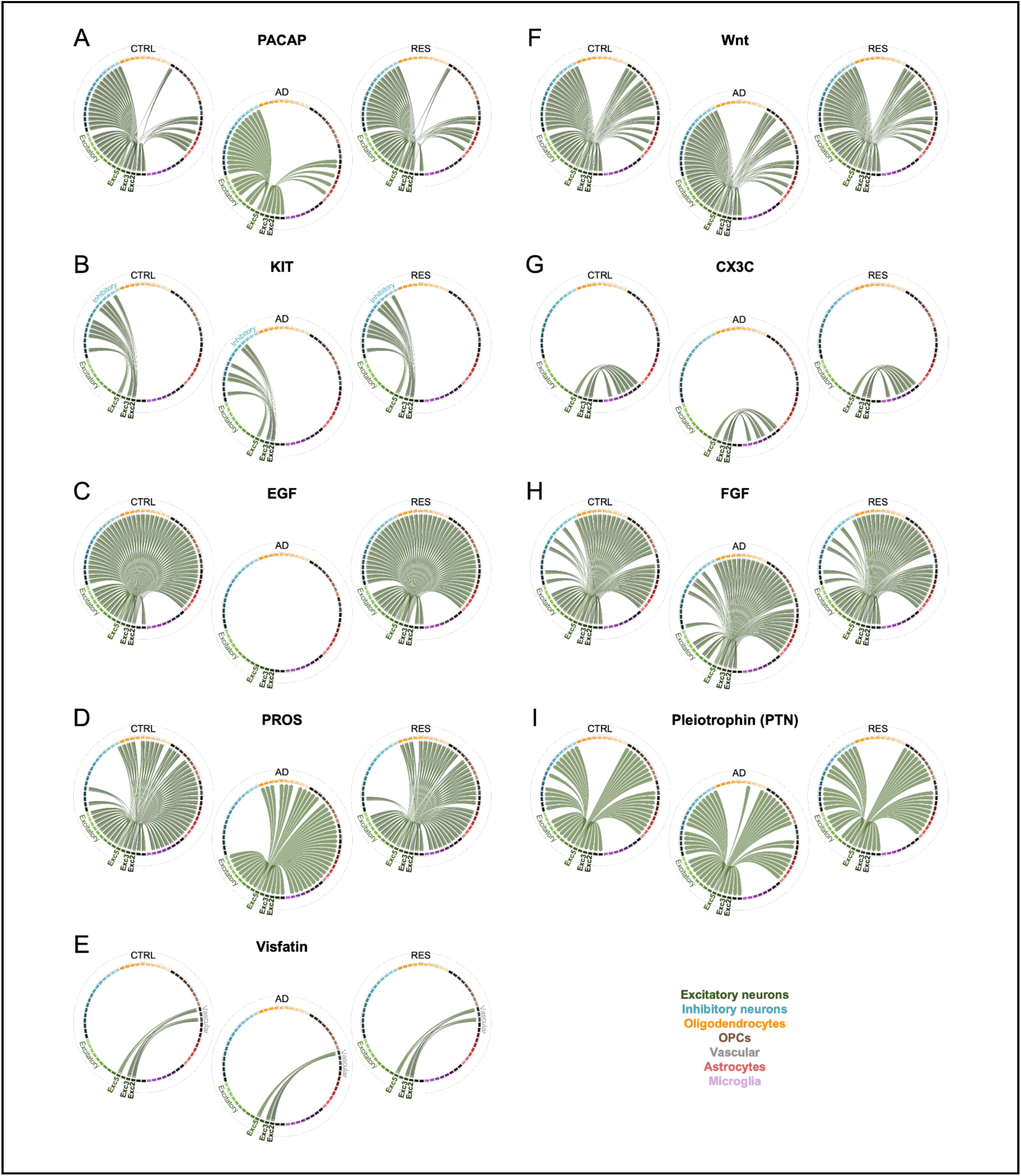
Signaling pathways dynamics in AD in resilience-associated excitatory neuronal subpopulations from the entorhinal cortex. Chord diagrams showing significant networks (Figure S15) with EC:Exc2, EC:Exc:3, and EC:Exc5 as sources. **(A)** PACAP (sources: EC:Exc2 and EC:Exc5; targets: subtypes of multiple major cell types; ligand: ADCYAP1; receptor: ADCYAP1R1). **(B)** KIT (sources: EC:Exc2, EC:Exc:3, and EC:Exc5; target: EC:Inh6, EC:Inh10, and EC:Inh14; ligand: KITLG; receptor: KIT). **(C)** EGF (source: EC:Exc2 and EC:Exc5; targets: subtypes of all major cell types except microglia; ligands: EGF and BTC; receptors: EGFR and ERBB4). **(D)** PROS (source: EC:Exc2; targets: subtypes from all major cell types; ligand: PROS1; receptors: AXL, TYRO3, MERTK). **(E)** Visfatin (sources: EC:Exc2, EC:Exc3, EC:Exc5; target: EC:Fib; ligand: NAMPT; receptor:ITGA5/ITGB1). **(F)** Wnt (sources: EC:Exc2, EC:Exc3, EC:Exc5; targets: EC:Opc4 (disappears in AD), and EC:Exc14 (emerges in AD); ligands: WNT10B, and WNT3 (in AD only); receptor: FZD3/LRP6). **(G)** CX3C (sources: EC:Exc2, EC:Exc3, EC:Exc5; target: EC:Mic2; ligand: CX3CL1; receptor: CX3CR1). **(H)** FGF (sources: EC:Exc2, EC:Exc3, EC:Exc5; targets: multiple subtypes of excitatory neurons, inhibitory neurons (loss in some subtypes and gain in others), and EC:Vas0 (emerges in AD); ligands: FGF5, FGF9, and FGF17; receptors: FGFR1 and FGFR2). **(I)** Pleiotrophin (source: EC:Exc5; targets: EC:Exc13 (gain), EC:Inh6 and EC:Inh8 (gain), EC:Oli6 and EC:Oli9 (gain), EC:Fib (loss), and EC:Ast2 (loss); ligand: PTN; receptors: PTPRZ1, SDC2 (loss), SDC3, NCL, ALK). CTRL: Control, AD: Alzheimer’s disease, RES: Resilient.

**Figure S17.**
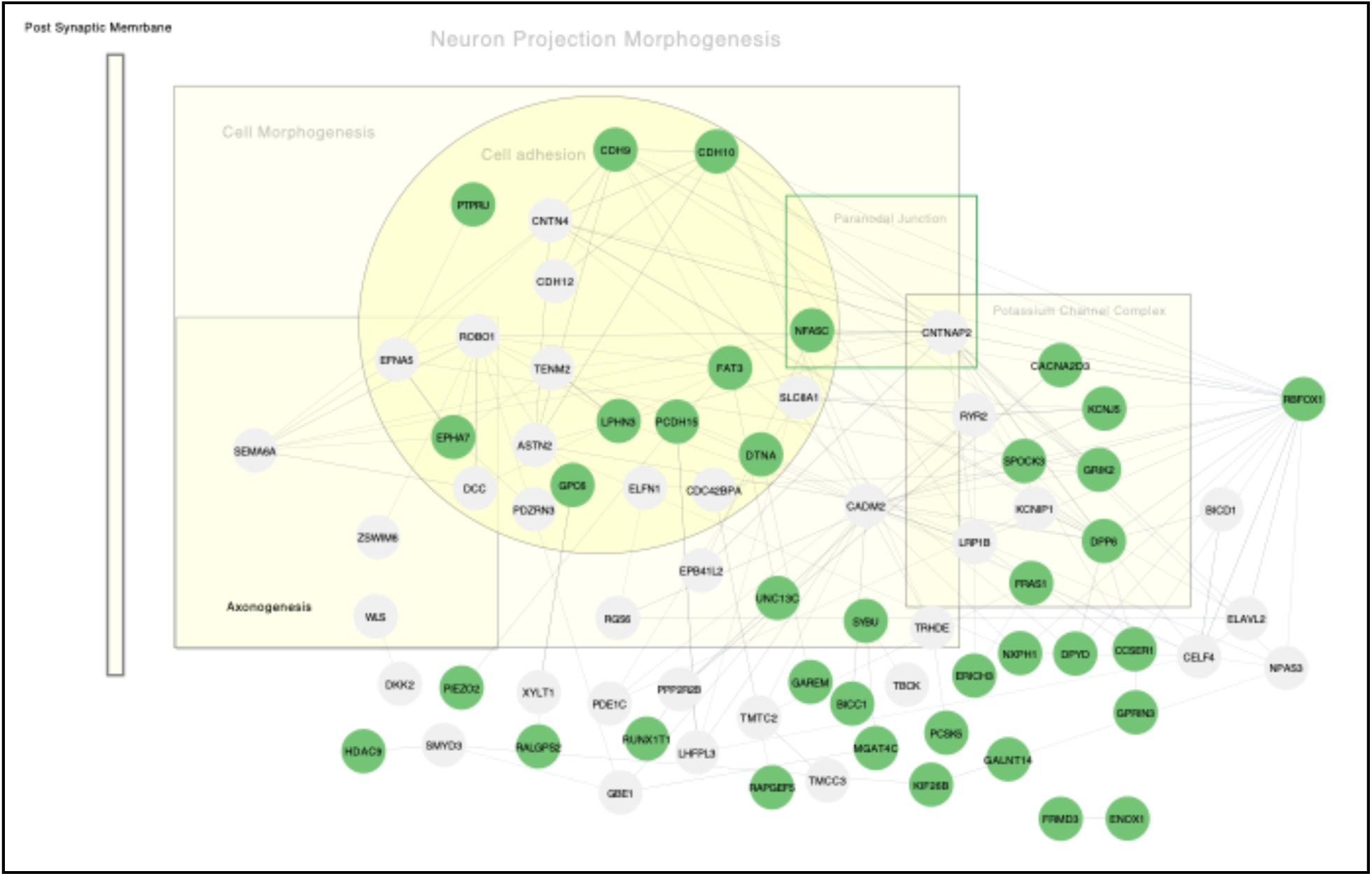
StringDB network of genes associated with risk and protective rare variants also identified as marker genes for DLPFC:Inh1 neurons. RBFOX1 interaction partners and marker genes are enriched in cell morphogenesis, paranodal junction, potassium channel complex, and axonogenesis. All may contribute to neuron projection morphogenesis. Genes from protective variants are shown as green nodes, and genes from risk variants as gray nodes. Visualization was performed using Cytoscape 3.10.2. (NDEx access to network).

**Figure S18.**
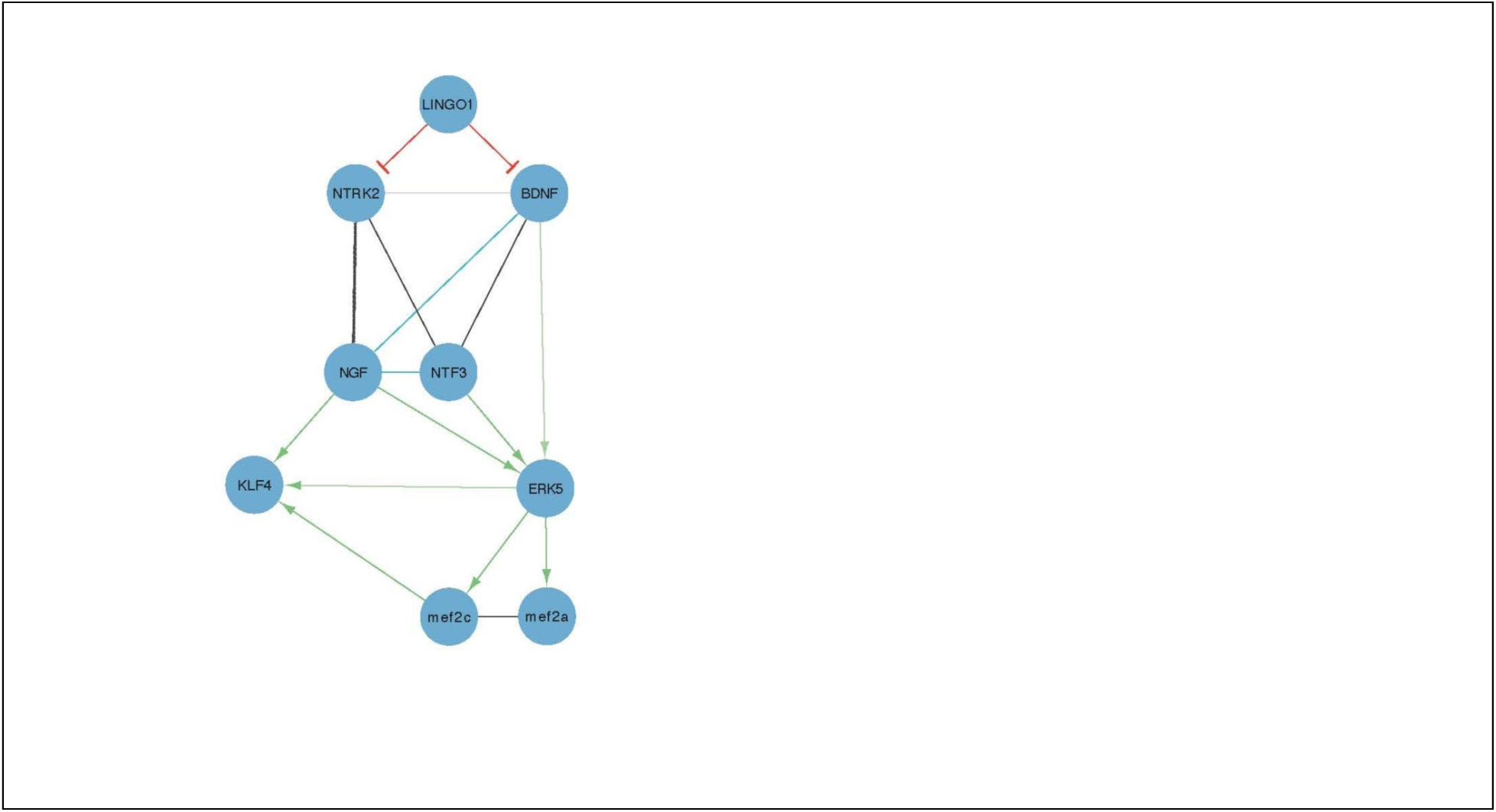
LINGO1 regulation of neurotrophin pathway in resilient cells. STRINGdb relationships and supporting literature were used to establish protein-protein and coexpression relationships between key members of the neurotrophin pathway. Visualization was performed using Cytoscape 3.10.2. Red-inhibition, green arrow, activation. Black, interaction, gray, imputed interaction.

## Supplementary tables

**Table S1. Differentially expressed genes in AD compared to cognitive resilience in bulk DLPFC tissue.**

Differential gene expression analysis was performed between AD and resilient subjects (ADvsRES) using ROSMAP bulk RNAseq data. Significant differential expression was determined using adjusted p-value (FDR) < 0.1 (highlighted in red) and |FC| > 1.1 criteria. P-values were determined using a linear regression model accounting for confounding covariates such as sequencing batch, RNA integrity number, and postmortem interval. The column “Direction” shows significant upregulation (“UP”) or downregulation (“DOWN”) in AD versus resilience as defined above. The column “Unique ADvsRES” reports genes that were differentially expressed in ADvsRES and not in ADvsCTRL or RESvsCTRL. The column “Cognitive loss” reports genes also associated with loss of cognition (related with **Table S5**).

**Table S2. Differentially expressed genes in AD compared to cognitively healthy controls in bulk DLPFC tissue.**

Differential gene expression analysis was performed between AD and control subjects (ADvsCTRL) using ROSMAP bulk RNAseq data. Significant differential expression was determined using adjusted p-value (FDR) < 0.1 (highlighted in red) and |FC| > 1.1 criteria. P-values were determined using a linear regression model accounting for confounding covariates such as sequencing batch, RNA integrity number, and postmortem interval. The column “Direction” shows significant upregulation (“UP”) or downregulation (“DOWN”) in AD versus controls as defined above.

**Table S3. Differentially expressed genes in resilient subjects compared to cognitively healthy controls in bulk DLPFC tissue.**

Differential gene expression analysis was performed between resilient and control subjects (RESvsCTRL) using ROSMAP bulk RNAseq data. Significant differential expression was determined using adjusted p-value (FDR) < 0.1 (highlighted in red) and |FC| > 1.1 criteria. P-values were determined using a linear regression model accounting for confounding covariates such as sequencing batch, RNA integrity number, and postmortem interval. The column “Direction” shows significant upregulation (“UP”) or downregulation (“DOWN”) in resilience versus controls as defined above.

**Table S4. Differentially expressed genes in AD compared to subjects classified as presymptomatic in bulk DLPFC tissue.**

Differential gene expression analysis was performed between AD and presymptomatic subjects (ADvsPRE) using ROSMAP bulk RNAseq data. Significant differential expression was determined using adjusted p-value (FDR) < 0.1 (highlighted in red) and |FC| > 1.1 criteria. P-values were determined using a linear regression model accounting for confounding covariates such as sequencing batch, RNA integrity number, and postmortem interval. The column “Direction” shows significant upregulation (“UP”) or downregulation (“DOWN”) in AD versus presymptomatic as defined above.

**Table S5. Genes associated with loss of cognition from bulk RNAseq data.**

The association of gene expression profiles with loss of cognition was analyzed using a proportional odds model (POM) for ordinal categorical data analysis applied to ROSMAP bulk RNAseq data. Gene expression levels were adjusted for confounding covariates prior to performing POM. The POM analysis was applied to the subjects with either no cognitive impairment, mild cognitive impairment, or AD dementia. The POM analyzed cognitive impairment as a function of gene expression and pathology status (plaque and tangle stages). P-values represent the significance of the association of each gene with cognitive status. The column “Direction” denotes whether the expression of a gene positively or negatively correlates with loss of cognition. The column “ADvsRES” denotes genes that were also differentially expressed between AD and resilient subjects (related with **Table S1**).

**Table S6. Dysregulated pathways in AD compared to cognitive resilience in bulk DLPFC.**

Differential pathway activity analysis was performed between AD and resilient subjects based on ROSMAP bulk RNAseq data. Significant pathway dysregulation was determined using Storey-adjusted p-value (q-value) < 0.1. P-values were determined using a linear regression model that accounts for confounding covariates such as sequencing batch, RNA integrity number, and postmortem interval.

**Table S7. Unsupervised clusters of dysregulated pathways in AD compared to cognitive resilience from bulk DLPFC.**

Cluster membership for each of the 99 dysregulated pathways in ADvsRES (related to **Table S6**) following unsupervised clustering. Modules of dysregulated pathways in ADvsRES were determined by mapping to the pathway co-expression network followed by the Label Propagation clustering algorithm. The column “cluster” represents the unsupervised clusters assigned by the described analysis. The analysis was performed using the PanomiR package with default parameters.

**Table S8. Dysregulated pathways in AD compared to cognitively healthy controls in bulk DLPFC.**

Differential pathway activity analysis was performed between AD and control subjects based on ROSMAP bulk RNAseq data. Significant pathway dysregulation was determined using Storey-adjusted p-value (q-value) < 0.1. p-values were determined using a linear regression model accounting for confounding covariates such as sequencing batch, RNA integrity number, and postmortem interval.

**Table S9. Results for the pathway activity analysis in resilient individuals compared to subjects classified as presymptomatic in bulk DLPFC.**

Differential pathway activity analysis was performed between RES and presymptomatic subjects (RESvs PRE) based on ROSMAP bulk RNAseq data. Significant pathway dysregulation was determined using Storey-adjusted p-value (q-value) < 0.1. P-values were determined using a linear regression model that accounts for confounding covariates such as sequencing batch, RNA integrity number, and postmortem interval.

**Table S10. Results for the pathway activity analysis in AD compared to presymptomatic subjects in bulk DLPFC tissue.**

Differential pathway activity analysis was performed between AD and presymptomatic subjects (ADvs PRE) based on ROSMAP bulk RNAseq data. Significant pathway dysregulation was determined using Storey-adjusted p-value (q-value) < 0.1. P-values were determined using a linear regression model that accounts for confounding covariates such as sequencing batch, RNA integrity number, and postmortem interval.

**Table S11. Number of subjects and number of cells per group defined in this study for snRNAseq data.**

**Table S12. Distributions for differentially expressed genes per comparison, identified from snRNAseq data for each major cell type in each brain region investigated.**

Differential gene expression analyses were performed per major cell type for each brain region by implementing a statistical model by group using the MAST statistical framework in *Seurat*, after removing non-variable genes. P-values were adjusted using the Bonferroni correction, as recommended for the R package *Seurat*. Significant differential expression was determined using adjusted p-value (adj-P) < 0.1 and log2FC > 0.2 or log2FC < −0.2 (|FC| > 1.1) criteria. The number of down-regulated differentially expressed genes (DEGs) in the first group compared to the second group are highlighted in red, and the number of up-regulated DEGs in the first group compared to the second group are highlighted in blue.

**Table S13. Selected differentially expressed genes for major cell types.**

Differential expression summary results for each brain region across the three comparisons (ADvsRES, ADvsCTRL, and RESvsCTRL) for genes discussed in the text. Differential gene expression analyses were performed per major cell type for each brain region by implementing a statistical model by group using the MAST statistical framework in *Seurat*, after removing non-variable genes. P-values were adjusted using the Bonferroni correction (adj-P), as recommended for the R package *Seurat*. Significant differential expression was determined using adjusted adj-p < 0.1 and log2FC > 0.2 or log2FC < −0.2 (|FC| > 1.1) criteria. NS: not significant. Direction “NONE” refers to log2FC below our threshold.

**Table S14. Cell annotations for cell subtypes for each brain region.**

Cluster annotations were generated using the web tool MapMyCells using the Hierarchical algorithm.

**Table S15. Results for cell proportion analysis per major cell types for each brain region investigated.**

Changes in cell composition between groups for each cell subpopulation (subclusters) were detected using a Dirichlet multinomial regression model, while accounting for the proportions of all of the other cell subsets within each major cell type. P-values were adjusted for multiple comparisons using the Benjamini-Hochberg (FDR) correction (adj-P). Green: Adj-P < 0.05, yellow: adj-P < 0.01, red: adj-p < 0.001.

**Table S16. Phenotypic and clinical characteristics of the BIDMC cohort.**

Human brain tissue used for immunostaining experiments was collected at Beth Israel Deaconess Medical Center (BIDMC) upon autopsy. Samples were tested for multiple pathologies, including TDP-43. Donors presenting comorbidities, including diabetes, were excluded.

**Table S17. Ligand-receptor results from CellChat for the SST signaling pathway in the EC.**

Source cell subtypes included EC:Inh3 and EC:Inh9 in control and resilient subjects, but in AD only EC:Inh3 and EC:Inh11 were identified as sources for the SST pathway in AD. The receptor SSTR1 was only observed in resilience.

**Table S18. Selected differentially expressed genes for cell subtypes.**

Differential expression summary results for each brain region across the three comparisons (ADvsRES, ADvsCTRL, and RESvsCTRL) for genes discussed in the text. Differential gene expression analyses were performed per cell subtype for each brain region by implementing a statistical model by group using the MAST statistical framework in *Seurat*, after removing non-variable genes. P-values were adjusted using the Bonferroni correction (adj-P), as recommended for the R package *Seurat*. Significant differential expression was determined using adjusted adj-p < 0.1 and log2FC > 0.2 or log2FC < −0.2 (|FC| > 1.1) criteria. NS: not significant. Direction “NONE” refers to log2FC below our threshold.

**Table S19. Interaction partners of RBFOX1 identified as rare variant-associated genes and marker genes for DLPFC:Inh1 neurons**

**Table S20. Functional enrichment of RBFOX1 partners, also identified as rare variant associated genes and marker genes for DLPFC:Inh1 neurons.**

## Abbreviations

AD: Alzheimer’s Disease
AD-PRS: Alzheimer’s disease polygenic risk scores
adj-P: Adjusted P-value
ADvsCTRL: Alzheimer’s disease versus control
ADvsPRE: Alzheimer’s disease versus presymptomatic
ADvsRES: Alzheimer’s disease versus resilient
ANGPT: Angiopoietin
ANGPT2: Angiopoietin-2
APOE: Apolipoprotein E
APP: Amyloid precursor protein
ATP8B1: ATPase phospholipid transporting 8B1
Aβ: Amyloid beta
BDNF: Brain-derived neurotrophic factor
BMP: Bone morphogenetic protein
BTC: Betacellulin
CA1: Cornu Ammonis 1
cGAS: Cyclic GMP-AMP synthase
CTRL: Control
DE: Differential expression
DEGs: Differentially expressed genes
DLPFC: Dorsolateral prefrontal cortex
EC: Entorhinal cortex
EC:Exc2: Excitatory neurons in the entorhinal cortex subtype 2
EC:Exc3: Excitatory neurons in the entorhinal cortex subtype 3
EGF: Epidermal growth factor
EGFR: Epidermal growth factor receptor
ERBB4: Erb-B2 receptor tyrosine kinase 4
ERK: Extracellular signal-regulated protein kinase
EWCE: Expression Weighted Celltype Enrichment
FC: Fold change
FDR: False discovery rate
FFPE: Formalin-fixed paraffin-embedded
GABA: Gamma-aminobutyric acid
GFAP: Glial fibrillary acidic protein
GO: Gene ontology
GWAS: Genome-wide association study
HC: Hippocampus
Hsp110: Heat shock protein 110
Hsp40: Heat shock protein 40
Hsp70: Heat shock protein 70
Hsp90: Heat shock protein 90
HSP90AA: Heat shock protein 90 alpha family class A
HSP90AA1: Heat shock protein 90 alpha family member A1
HSP90AB1: Heat shock protein 90 alpha family class B member 1
Hsps: Heat shock proteins
IFN-I: Interferon type I
IL-1β: Interleukin 1 beta
IL-6: Interleukin 6
ITGA5: Integrin alpha-5
ITGB1: Integrin beta-1
KIF26B: Kinesin family member 26B
KIT: KIT Proto-oncogene
KLF4: Kruppel-like factor 4
KLF4: Krüppel-like factor 4
L-R: Ligand-receptor
LCM: Laser capture microdissection
LINGO1: Leucine-rich repeat and Ig domain-containing protein 1
Log2FC: Log2 fold change
MAPK: Mitogen-activated protein kinase
MEF2A: Myocyte enhancer factor 2A
MEF2C: Myocyte enhancer factor 2C
mIF: Multiplex immunofluorescence
MSigDB: Molecular Signatures Database
ncWnt: Non-canonical Wnt
NGFR: Nerve growth factor receptor
NRN1: Neuretin
NRTK: Neurotrophin receptor tyrosine kinase
NT: Neurotrophin
NTRK2: Neurotrophic receptor tyrosine kinase 2
OPCs: Oligodendrocyte progenitor cells
PCxN: Pathway Co-expression Network
PI3K: Phosphoinositide 3-kinase
PLCγ: Phospholipase Cγ
PRE: Presymptomatic
PREvsCTRL: Presymptomatic versus control
PRS: Polygenic risk score
PS1: Presenilin 1
pTau: Phosphorylated tau
PVALB: Parvalbumin
RBFOX1: RNA Binding fox-1 Homolog 1
RELN: Reelin
RES: Resilient
RESvsCTRL: Resilient versus control
RESvsPRE: Resilient versus presymptomatic
ROSMAP: Religious Order Study and Memory and Aging Project
SNPs: Single nucleotide polymorphisms
snRNAseq: single-nucleus RNA sequencing
SORT1: Sortilin 1
SST: Somatostatin
SSTR1: Somatostatin receptor 1
SSTR2: Somatostatin receptor 2
TEK: TEK receptor tyrosine kinase
TGF-β: Transforming growth factor beta
TREM2: Triggering receptor expressed on myeloid cells 2
TULP: Tubby-like proteins
UMAP: Uniform Manifold Approximation and Projection
XBP1s: X-box binding protein 1 spliced

